# Characterizing adjuvants’ effects at the murine immunoglobulin repertoire level

**DOI:** 10.1101/2022.11.19.517218

**Authors:** Feng Feng, Rachel Yuen, Yumei Wang, Axin Hua, Thomas B. Kepler, Lee Wetzler

## Abstract

High-throughput immunoglobulin sequencing (IgSeq) has been developed and applied to study the adaptive immune response extensively for more than a decade. However, generating large-scale, high-fidelity sequencing data is still challenging, and furthermore, not much has been done to characterize adjuvants’ effects at the repertoire level. Thus, we developed an improved library prep protocol and standardized the data analysis pipeline for accurate repertoire profiling. In addition, two metrics were implemented to assess repertoire clone properties. We then studied systemically the effects of two adjuvants, CpG and Alum, on the Ig heavy chain repertoire using the ovalbumin (OVA) challenged mouse model. Ig repertoires of different tissues (spleen and bone marrow) and isotypes (IgG and IgM) were examined and compared in terms of sequence mutation frequency, IGHV gene usage, CDR3 length, rescaled Hill numbers for clonal diversity, and clone selection strength. As a result, Ig repertoires of different tissues or isotypes exhibited distinguishable profiles at the non-immunized steady state. Adjuvanted immunizations further resulted in statistically significant alterations in Ig repertoire compared with PBS or OVA alone immunized groups. Lastly, we applied unsupervised machine learning techniques – multiple factor analysis and clustering – to identify Ig repertoire signatures in different compartments and under varying immunizations. We found that the IGH repertoires of distinct tissue-isotype compartments or under varying immunizations differed in unique sets of properties. Notably, Alum and CpG effects on the Ig repertoire exhibited different tissue and isotype preferences. The former led to increased diversity of abundant clones of both isotypes in BM only, and the latter promoted the selection of IgG clones only but in both tissues. The patterns of Ig repertoire changes likely reflected possible action mechanisms of these two adjuvants.

## Introduction

With the advent of technological advancements in the last two decades, immunoglobulin sequencing (IgSeq), allowing for sampling immunoglobulin variable region genes (IgVRG) at an unprecedented depth, has proved invaluable to study adaptive immune responses in health^1^ and disease^2, 3^, and for diagnostic^4^ and therapeutic^5^ purposes. However, generating large-scale, high-fidelity, and high-quality sequencing data is challenging^6, 7^; there is room for further improvement in methods for both data generation and data analysis^8, 9^.

Adjuvants are critical in almost all vaccines to increase antigen immunogenicity and enhance the magnitude and durability of the response. Despite their extensive use in vaccines for more than a century, the understanding of adjuvants’ action mechanisms remains partial^10^. As a result, only very few adjuvants have been licensed for human use. Among them are aluminum (Alum) salts, the first and most commonly used adjuvant. Alum adjuvants are particulates and induce antibody and CD4+ T helper cell responses^10^. It has long been accepted that Alum exerts its effects through creating an *in situ* depot, which as a delivery system allows for slow release of antigens from the site of immunization and enhanced antigen uptake by DCs^11^. However, recent reports suggested that a short-term depot was not necessary for the effect of aluminum adjuvants^12^. Alum also, potentially, enhances the immune response through several other mechanisms. It induces tissue release uric acid, which activates the intracellular NLRP3 inflammasome and increases pro-inflammatory IL-1β secretion^10, 11^. However, the data are contradictory regarding the importance of the NLRP3 pathway in Alum’s adjuvant effect^10, 11^. Moreover, Alum can increases recruitments of various immune cells, including DCs, natural killer cells, and IL-4 secreting eosinophils which in turn promote the Th2 responses commonly seen with the use of Alum ^10, 13^. Unmethylated deoxycytidyl-deoxyguanosine (CpG) oligonucleotide ligands are another group of adjuvants used in licensed vaccines. By mimicking bacterial DNA signatures, CpG triggers cells expressing the toll-like receptor 9 (TLR9). There are at least four different types of CpG, distinguishable by their nucleotide sequences and effects on immune responses^14^. Yet, they all influence many immune cells, especially DCs and macrophages. CpG enhances pDC migration and activates them to produce type 1-IFN plus a few other cytokines. It also boosts B cell differentiation, proliferation, and antibody production (such as IgM)^14, 15^. Lastly, CpG directly or indirectly stimulates NK cells and Th1 CD4+ cells, resulting in NK activation and an improved antigen specific CD4 and CD8+ T cell response^10, 14, 15^. Again, despite these recent insights, the immunological mechanism and characterization of adjuvant effects is still incomplete. Furthermore, the adjuvant effects on the immune repertoire are not well studied.

B cells in different tissues exhibit great functional and phenotypic heterogeneity. The spleen, the largest secondary lymphoid organ, plays a critical role in bridging the innate and adaptive immunity^16^, and makes up in a large portion the peripheral naïve B cell pool, including transition B cells, follicular B cells, marginal-zone B cells^17^. It is also a major hub for long-lived memory B cells^18^ and possibly some antibody-secreting plasma cells, among which are long-lived^17^, in the spleen red pulp. On the other hand, bone marrow (BM) is both a primary lymphoid organ for B cell development and maturation and also a vital long-term niche for terminally-differentiated plasma cells^19, 20^ plus other types of immune cells. Such a specialized niche is not static but dynamically responds to physiological and pathological challenges and, in turn, regulates B cell lymphopoiesis and plasma cell survival/turnover ^19, 20, 21, 22^. In addition, B cells of different isotypes also represent heterogeneous cell subsets with distinct intrinsic properties in terms of their location, origin, developmental stage, differential potential, and effector function. Recent work has reported changes in the IGH repertoire of B cells of different isotypes with metrics such as gene usage, mutation, and clonality^2, 23, 24^. Thus, it would be wise to assume B cells of varying tissues and isotypes to form distinct compartments.

The B cell Ig repertoire has, by some estimates, the theoretical potential to comprise up to 10^14^ unique molecules in humans^25^ and 10^13^ in mice^26^. Such a diverse repertoire is essential to achieve immunity against possible encountered antigens during one’s lifetime. Traditionally, B cell responses were evaluated by assessing the quantity and function of one or a few specific antibodies in individuals. Unfortunately, those measures reflect little about changes at the B cell repertoire level. Characterization of total repertoire changes has proven insightful for understanding vaccine responses^2, 3, 24^. In this work, we immunized mice with ovalbumin (OVA) mixed with either Alum or CpG to systemically study the effects of these adjuvants on the Ig repertoire. We developed an in-house Ig heavy chain library prep protocol and data processing pipeline to generate high-fidelity and high-quality IgSeq data. Heavy-chain repertoires were profiled for four different tissue-isotype compartments from mice under varying immunization conditions (PBS, OVA alone, OVA+CpG, OVA+Alum). We examined sequence mutation frequency, IGHV usage, and CDR3 length of these Ig repertoires. In addition, we developed two metrics to characterize the Ig repertoire at the clone level, rescaled Hill numbers for clone diversity and clone selection strength. We found that Ig repertoires showed statistically significant alterations in these properties among different compartments at the unimmunized steady state and further diverged after immunization. Finally, we characterized the Ig repertoire signatures of different tissues, isotypes, and adjuvant immunizations by unsupervised machine learning approaches. We found that OVA immunization with alum or CpG gave rise to repertoires distinct from each other and from those under immunization with unadjuvanted OVA and displaying different tissue and isotype preferences. Our data suggested possible action mechanisms of adjuvants: Alum resulted in increased diversity of abundant clones in BM repertoires of both isotypes; CpG led to the increased selection of IgG clones and affected BM and spleen compartments similarly.

## Methods and Materials

### Mouse immunization

C57BL/6J mice were purchased from The Jackson Laboratory (Bar Harbor, ME, USA). At the age of 6-8 weeks, each group of three mice started to receive one of the four subcutaneous immunizations in a volume of 100μl with 10μg ovalbumin (OVA) plus adjuvant or vehicle control (PBS): PBS, OVA, OVA + CpG (10 μg, Invivogen, Cat# ODN1826) and OVA + Alum (200 μg Sigma, Cat# A8222). A total of four immunizations were applied at two-week intervals between the first three injections and a four-week interval between the third and the fourth dose (Fig. 1A). All mice were maintained in the Association for Assessment and Accreditation of Laboratory Animal Care International accredited facility at Boston University School of Medicine Laboratory Animal Science Center. All animal experiments were conducted under the approved Institutional Animal Care and Use Committee (IACUC) protocol at Boston University.

**Figure 1.**
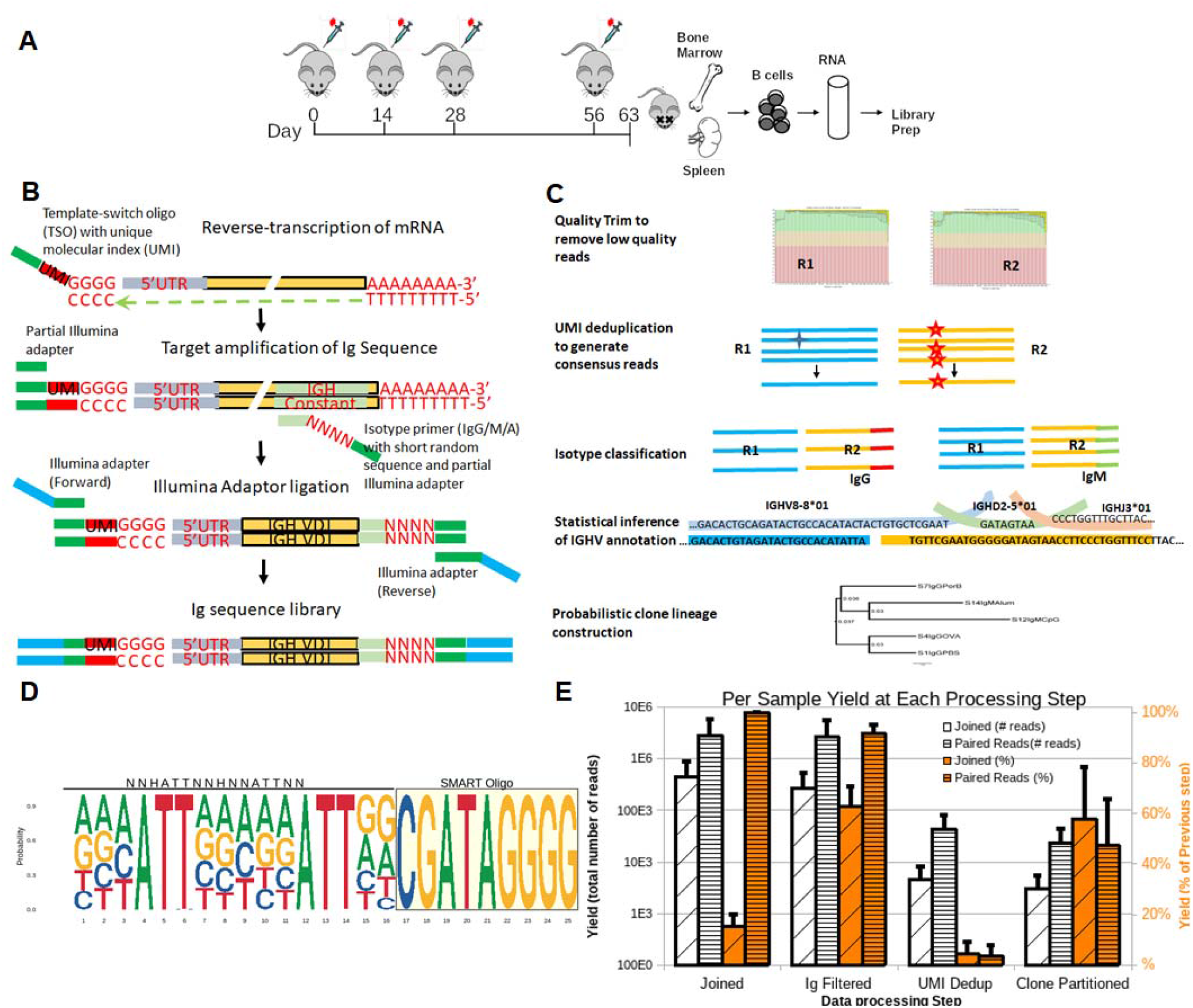
Sequencing library preparation protocol and data preprocessing pipeline. A) Each group of three mice received four doses of immunizations (PBS, OVA, OVA+CpG and OVA+Alum). One week after the fourth injection, spleen and bone marrow were harvested. Splenic B cells and bone marrow cells were prepared and total RNA was isolated. B) The 5’RACE library prep protocol started with the TSOs containing 5’-biotin blockers and UMIs including spacers and RT-PCR cycles running at a higher temperature. The target Ig amplification PCR was achieved with a reverse isotype-specific primer containing short random nucleotide to increase the sequence diversity in the beginning of Read 2 sequencing. Both PCRs were monitored at real time to avoid over-amplification. Pair end sequencing was done on an Illumina HiSeq2500 system. C) The data preprocessing pipeline does not require pair-read joining and in all steps separated reads were used as input. We also developed and implemented new algorithm to de-duplicate UMI group reads and generate consensus sequences. Ig heavy chain VDJ annotation and clonal analysis were carried out using the Cloanalyst software. D) The sequence logo plot for the TSO with UMIs. The UMI is different from a typical one by including spacers with reduced degeneracy. E) Sequence yield of each sample at different steps between the pair-end joining protocol and non-joining pipeline in this work. Open bars are for the sequence yields of the typical pair-joining protocol and filled bars are for the non-joining separated reads pipeline. White bars are yields in number of sequences and orange yields in percentages. In the end, with the non-joining pipeline on average we obtained about 10-fold more sequences and 8-fold more clones for each sample.

### Cell collection and IGH sequencing library prep

A week after the fourth immunization, BMs and spleens were harvested. The spleen was disrupted, and cell suspension was obtained by passing a 70μm mesh nylon strainer. Splenic B cells were enriched with the EasySep kit (STEMCELL Technologies, Cambridge, MA). Bone marrow cells were obtained by flushing femora and tibiae with RPMI medium, and whole tissue cells were used without enrichment. Red blood cells were lysed from samples with ACK (150 mM NaH_4_CL, 50mM KHCO_3_) lysis buffer. Splenic B cells and bone marrow tissue cells were treated with RNAzol RT (Sigma-Aldrich), and total RNA was isolated according to the manufacturer’s instruction and saved at -80°C.

We adopted a 5’ rapid amplification of cDNA ends (5’RACE) approach for library prep (Fig. 1B) ^27, 28, 29, 30, 31, 32, 33^. cDNA synthesis was done with two primers, forward template switching oligos (TSO) with unique molecular identifiers (UMI) /5Biosg/-GTCAGATGTGTATAAGAGACAGNNHATTNNHNNATTNNCGATAGrGrGrG-3’ and reverse 5’- GTGTCACGTACAGAGTCATC(T)_30_VN-3’. Two hundred nanograms of total RNA was reverse-transcribed in the presence of 100uM primers, 10mM dNTP mix, and 40U/uL Maxima H Minus transcriptase (Thermo Scientific). PCR was run on a thermal cycler with a cycle: 30 minutes at 50°C for the initial step; 2 minutes at 50°C and 2 minutes at 42°C for 10 cycles; 15 minutes at 80°C to inactivate the transcriptase.

cDNA was cleaned with AmpureXP beads (Beckman Coulter Life Sciences, Indianapolis, IN) and amplified in the presence of 0.4uM primer mix, forward primer:5’- TCGTCGGCAGCGTCAGATGTGTATAAGAGACAG-3’ and reverse isotype-specific (IgG1/2:5’- GTCTCGTGGGCTCGGAGATGTGTATAAGAGACAGNNNNCAGATCCAGGGGCCAGTGGATAG A*C-3’; IgG3: 5’- GTCTCGTGGGCTCGGAGATGTGTATAAGAGACAGNNNNTGCAGCCAGGGACCAAGGGATAG A*C; IgM:5’- GTCTCGTGGGCTCGGAGATGTGTATAAGAGACAGNNNNGGGAAGACATTTGGGAAGGACTG A*C-3’). Then the last PCR was carried out to add Illumina adapters and dual multiplexing indexes (10nts long) with 0.125uM primer pairs, forward: AATGATACGGCGACCACCGAGATCTACAC-[i5 index]-TCGTCGGCAGCGT*C and reverse: CAAGCAGAAGACGGCATACGAGAT-[i7 index]-GTCTCGTGGGCTCG*G (Integrated DNA Technologies, Iowa, USA). Both PCR amplifications were done with KAPA Real-time library Amp kit (Roche, Wilmington, MA) with a cycle condition: 45 seconds at 98°C for initial denaturation; 15 seconds at 98 °C, 30 seconds at 66°C and 60 seconds at 72°C for up to 22 cycles; 2 minutes at 72°C for the last extension. PCRs were monitored in real-time and terminated when reaching a defined threshold to avoid over-amplification. Final PCR products were cleaned up by beads, measured for DNA concentrations by a Qubit 4 fluorometer (ThermoFisher Scientific), and then multiple samples were pooled with equal amounts. Finally, the library was size-selected between 400∼1000bp by Blue Pippin (LABGENE Scientific, Switzerland) before being sequenced on Illumina HiSeq2500 with 2×250bp chemistry. Libraries were loaded at 6.0pM with 5% Phix control v3 (Illumina) added.

### Data preprocessing

We preprocessed the sequence data using procedures developed in-house, including sample demultiplexing, quality trimming, search filtering, UMI deduplication, isotype determination, recombination parameter identification, and clonal partitioning. (Fig. 1C). Sample demultiplexing was done using *bcl2fastq* (Illumina) with no mismatch allowed. Quality trimming was done using Trimmomatic with a sliding window of 4 nucleotides and a minimum quality score of 20^34^. We ran bowtie2 comparing the sequences to an Ig heavy-chain database to filter out non-IG genes^35, 36^ (see Supplementary Material for details). Isotype classification was carried out using an in-house developed C++ program with flexibility for mismatches and offset isotype sequences. To meet the specific needs of Ig sequencing data, we implemented an in-house algorithm for deduplicating UMIs in python. Sequences within each group have identical UMIs, share the first 25 nucleotides identically, have less than 5% mismatches, and have identical isotype. To increase the quantification accuracy a directional network-based method was adopted to account for sequencing errors in UMIs^37^. Consensus sequences were generated on the basis of the multiple alignments of sequences of each group and taking into consideration of sequence quality. Singleton sequences were excluded from the analysis. Finally estimation of recombination parameters and clonal partitioning were done using Cloanalyst^38^ (see details in the next section). All scripts and software tools are available at https://github.com/BULQI/IgSeq and https://github.com/BULQI/Cloanalyst

### Heavy chain variable region gene usage

The IGHV counts for each sample were converted to compositional data using the R compositions library^39^, and further transformed to center log-ratios (CLR) to map the Aitchison simplex into real space^39^. The principal component analysis (PCA) was performed using the R stats package. The coordinates in the first few PCs were used as covariates in multi-factor analysis of variance (ANOVA). *Post hoc* analyses were performed using false discovery rate.

### Mutation frequency and CDR3 length

Mutations were identified by comparing IgVRG sequences with their inferred IGHV genes through the end of framework region 4 only (excluding CDR3). The mean and standard deviation of mutation frequencies were recorded for each of forty-eight groups. The mean and standard deviation of CDR3 lengths (nucleotides) for each sample were calculated. We then ran the three-way ANOVA on mean mutation frequencies and CDR3 lengths to test the effects of three factors (tissue, isotype, and immunization) (see *Statistical Analysis* for details).

### Rescaled clone diversity with Hill numbers

Clonal diversity was quantified using Hill numbers^41^, estimated using the methods described in Chao, et al. work^40^. To further reduce bias in the estimates from finite sample size, we made comparisons by pseudorandomly subsampling each sample to the size of the smallest sample a total of 100 times each. The ‘SpadeR’ R package^41^ was used to estimate diversity profiles as Hill numbers of order q between 0 and 4^40^, which were further rescaled by the clonal richness and transformed by the arcsine square root function. This rescaling was intended for down weighting the effect of clone richness on the diversity measure but emphasized the relative clone abundance information (or the clone evenness). Finally, three-way ANOVA and post hoc analysis were performed to test for factor effects.

### Strength of selection

To assess the level of selection that occurred within a B cell clone during its development, we compare two statistical characters of the associated phylogenetic tree: pairwise sequence dissimilarity (PSD) and mean mutation frequency (MMF). Our expectation is that clones experiencing stronger selection will exhibit lower ratio of sequence dissimilarity to mean mutation frequency. We calculated the MMF and PSD of the forty largest clones in each sample and made scatter plots. For comparisons, we established a threshold ratio to define the percentage of selected clones in each sample (Fig S19). Finally, the percentages of clones with selection were transformed to compositional data and used for statistical analysis.

### Unsupervised machine learning

To identify and summarize the feature variations characteristic of different treatments, we carried out multiple factor analysis (MFA) and unsupervised clustering on the complete assembled dataset. For each sample, there were five quantitative variables: mutation frequency, IGHV gene usage (multicomponent), CDR3 length, Hill numbers (multicomponent), and clonal selection level; these were used in the MFA. In addition, there were three supplementary categorical variables: tissue, isotype, and treatment (immunization). We conducted MFA using the ‘FactorMineR’ package^42^ and visualized data using the ‘factoextra’ R package.^42^ Unsupervised clustering was carried out on the PCs generated by MFA. Three clustering algorithms were performed, agglomerative hierarchical clustering (HC), spectral clustering, and affinity propagation clustering (APC). HC was done using the base R library and visualized using the ‘dendroextra’ R package,^43^ spectral clustering was done using the ‘kernlab’ R package^44^, and APC was done using the ‘APCluster’ R package^45^. Results were visualized on a 2D figure projected from high dimensional data with the ‘umap’ R package^46^.

### Statistical analysis

We ran three-factor ANOVA to test for effects of tissue (spleen and bone marrow), isotype (IgM and IgG), and immunization (PBS, OVA, OVA+CpG, and OVA+Alum) on repertoire properties (mutation, gene usage, etc.). The data were transformed to obtain normality and homoscedasticity. For IGHV gene usage data, we transformed gene counts into compositional data^39^, which were further center-log ratio (CLR) transformed to map the Aitchison simplex into real space. Similarly, normalized clone diversity data were arcsine transformed, and the CLR transformation was done on the clonal selection data, which are percentages of clones under selection in each sample.

Both type II and III ANOVA tables were obtained by the R ‘car’ package^47^ because of unequal sample sizes of different groups. For tests yielding significant differences (p<0.05) in ANOVA, we ran *post hoc* analysis by the simple effect test with contrasts using false discovery rate (FDR) to control for the multiple comparison error. The follow-up analyses were done using the R ‘emmeans’ package^48^. All R code and scripts are available at https://github.com/BULQI/IgSeq.

## Results

### IGH sequencing library prep and data preprocessing pipeline

For library prep, we developed an adapted 5’ rapid amplification of cDNA ends (5’RACE) approach (Fig 1B) ^27, 28, 29, 30, 31, 32, 33^, which is simple, unbiased, and requires only minimal amounts of material^27^. We improved the protocol by incorporating features for high-quality and accurate repertoire sequencing. In the cDNA reverse transcription step, we added 5’ biotin blockers to the template-switching oligos (TSO) ^49^ and ran reactions with cycles at a higher temperature (50°C)^29^. The former was to reduce the background concatamer formation (“hedgehog effects”), and the latter was to promote the unfolding of RNA secondary structure for a better yield of cDNA products. Moreover, the TSO contained a unique molecular identifier (UMI) that can be used at the data analysis step to correct PCR bias^50^. The UMI contains spacers with either fixed nucleotides or a partial degeneracy (Fig. 1D). Such design aimed to reduce artifacts in library prep such as non-specific amplification, strand invasion, and primer-dimer^50, 51, 52, 53^. At the PCR amplification step, isotype-specific primers also contain at the 5-prime end degenerated nucleotides (7nts long), which in design increases the complexity of the read-2 sequences^54^. That is necessary since the read-2 sequences start at the junction between the IGHJ segment and the constant region, and both parts consist of low diversity/complexity sequences (Fig. S2). Increasing diversity at the beginning of the read-2 resulted in a significantly higher quality of sequencing (supplementary Fig. S1 and Table S1). Lastly, we incorporated the unique dual indexes (distinct pairings) for sample multiplexing to avoid the indexing hopping^55^. Both amplification and adaptor ligation PCRs were monitored in real-time and terminated when PCR products reached a defined threshold to avoid over-amplification.

We developed a new data preprocessing pipeline that can utilize the features incorporated in the library prep method to increase quantification accuracy (Fig. 1C). Typical IgSeq data analysis pipelines require the assembly of paired reads (pair-read joining) ^9, 56, 57^. Sequence lengths, however, vary depending on factors such as primer location, V(D)J segment usage and recombination, the inclusion of leader regions, etc^50, 58^. Also, the assembly of two reads typically requires an overlap between them. Filtering out reads that fail to overlap leads to a bias towards short Ig and CDR3 sequences (Fig. S3). Therefore, we developed a pipeline that does not require overlap-based joining. One immediate benefit was more usable reads obtained through the new pipeline (Fig 1E). On the same sequencing data set, pair-read joining excluded ∼85% of raw reads. In addition, pair-read joining relying on a short sequence overlap between paired reads might result in erroneous assembly. These assemblies resulted in a lower yield in the Ig filtering step (60% Ig genes for pair-joining data vs. 90% Ig genes for the new pipeline, Fig. 1E). After the UMI deduplication step, pair-end joining resulted in ∼5×10^3^ unique Ig sequences per sample, and the processing with no pair-end joining yielded ∼5×10^4^ unique Ig sequences each. Lastly, the new pipeline allowed us to identify, on average, about 2.3×10^3^ clones per sample, which was eight-fold more than the number of clones obtained with the pair-end joining processing (Fig. 1E).

In the second step of the pipeline, we filtered out the non-IG genes using bowtie2 with an Ig gene segment database. Libraries prepared with 5’RACE can result in non-specific amplification of non-Ig genes that may skew downstream analyses, this step eliminates these off-target reads. Next, we ran the UMI deduplication to correct sequencing and PCR errors/bias. Our deduplication method is specifically suited to IgSeq, as follows. Each UMI is extended to include the first 25 nucleotides of the sequence and UMIs are merged based on a directional network-based algorithm^37^; multiple sequence alignments are used to identify sequence mismatches within each UMI group; groups with more than 5% mismatches or belonging to different isotypes are split; UMI groups with one single sequence are discarded; the consensus sequence for each UMI group is generated taking quality scores into account. All these steps were designed to correct sequencing and PCR errors (such as index hopping, non-specific amplification, strand invasion, and primer-dimer^50, 51, 52, 53^) and increase quantification accuracy.

Our approach produced isotype-aware sequences. We implemented an in-house C++ program to separate different isotypes. It is alignment-based and allows mismatches. Recombination parameter estimation and clonal partitioning were done using our software, Cloanalyst, which uses explicit likelihood models for V(D)J recombination and somatic mutation^38^.

Overall, we prepared IGH repertoire libraries from spleens and bone marrows of 15 mice and then obtained 84 million raw paired reads on sequencing. These reads yielded 1.2 million deduplicated Ig sequences assigned to 667,000 clones, which belong to 60 samples of four different tissue-isotype compartments (spleen IgM, spleen IgG, BM IgM, and BM IgG).

### Adjuvants impacted IGHV mutation frequency in a tissue- and isotype-dependent fashion

Somatic hypermutation (SHM), the driver of affinity maturation, has been used as a surrogate for the amount of antigen stimulation that a B cell clone has received during development^8, 59^. Here, we determined the influence that adjuvants have on the level of somatic hypermutation in different tissue-isotype compartments following immunization with OVA with or without CpG or alum. At the steady state (PBS controls), we observed altered mutation frequencies in different isotype or tissue compartments (IgG vs. IgM or BM vs. spleen). The IgG compartments in both tissues exhibited higher mean mutation frequencies than the IgM compartment, although no statistical significance was achieved (p>0.05, Fig 2A). Such a trend has been seen before^60, 61, 62^. The linking between isotype and hypermutation was likely a result of a common mechanism involving activation-induced cytidine deaminase (AID)^63, 64^. Upon comparing tissue differences, BM sequences exhibited a significantly higher mean mutation frequency than splenic sequences independent of isotypes (Fig. 2B, p<0.005). We expected such changes since the spleen, including its marginal zone, is a major reservoir for naive and memory B cells^18, 65, 66^; BM provides a preferred niche for plasma cells derived from the late germinal center output and enriched for high-affinity and highly mutated Ig sequences^65, 67, 68^.

**Figure 2.**
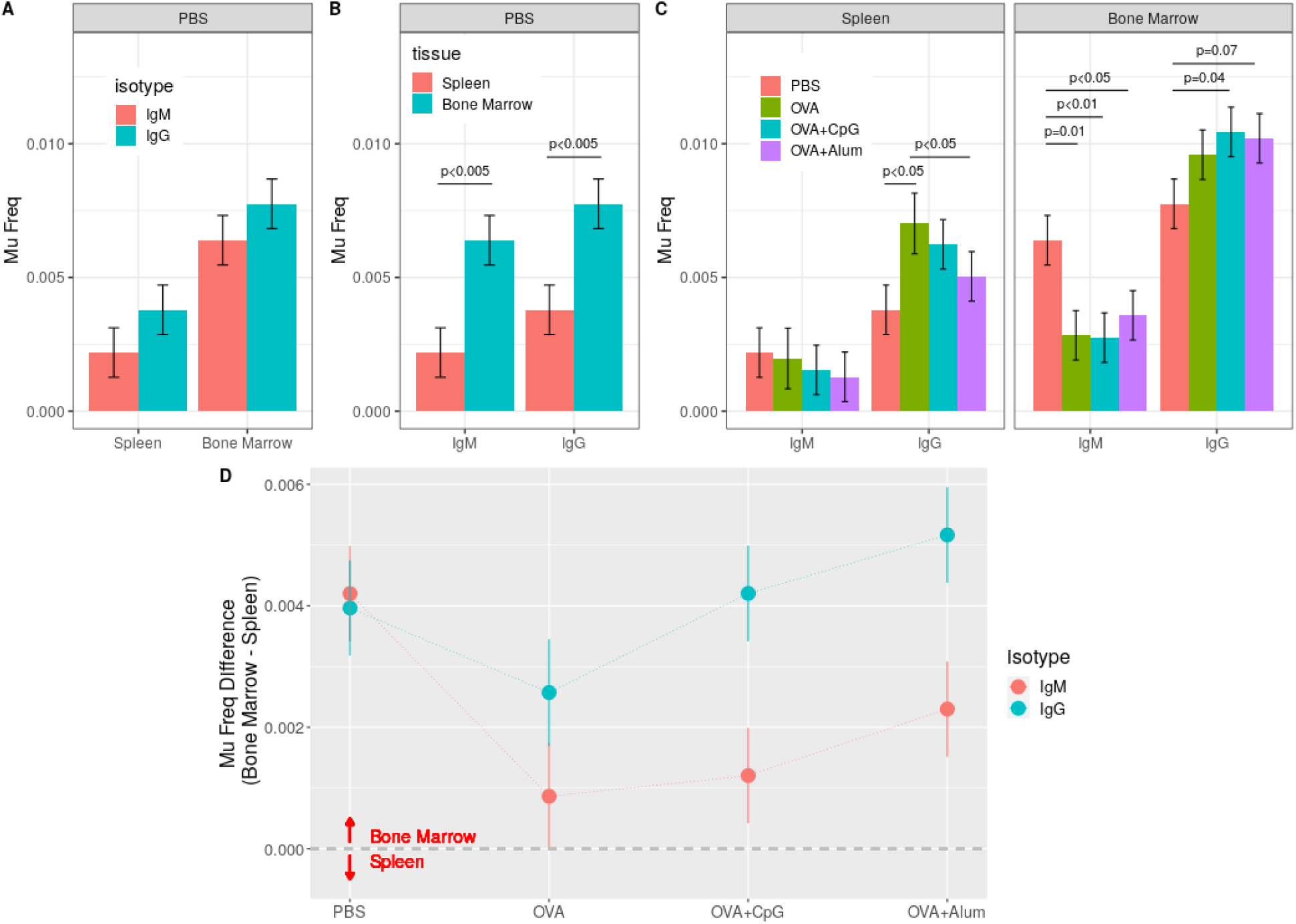
IGHV mean mutation frequency is affected by OVA immunizations and adjuvnats in a tissue- and isotype-dependent fashion. IGHV annotation was done by aligning to germline sequences with the software tool, Cloanalyst. Mutations were identified and expressed as per-nucleotide mutation frequency. The distributions of IGHV mutations and means and standard deviations were obtained and used for statistical tests. A) IGHV mutation frequencies in different isotype compartments in non-immunized mice (PBS controls). B) IGHV mutation frequencies in different tissue in non-immunized mice (PBS controls). C) IGHV mutation frequencies were affected by OVA immunization and the effects were isotype- and tissue-dependent. D) Mean mutation frequency difference between bone marrow and spleen. The difference was calculated as the BM mutation frequency subtracted by that of spleen for each OVA immunization group of different isotypes. A positive value means the BM compartment has a higher mutation frequency than the respective spleen compartment, a negative value means higher mutation frequency in the spleen, and zero means identical mutation frequency between two tissues. (Statistical p-values were corrected by FDR and significant p-values were indicated in figure)

Next, we observed that an OVA immunization (either alone or adjuvanted) increased the mean mutation frequency of IgG sequences but decreased that of IgM sequences compared with the PBS controls (Fig. 2C). This might suggest an effect of OVA immunization on promoting IgMs with high mutations to switch into IgGs. In addition, immunization effects further exhibited a tissue dependency (Fig. 2C). OVA alone yielded a higher mean mutation frequency in the spleen IgG compartment (p<0.05 for both OVA vs. PBS and OVA vs. OVA+Alum), while OVA immunizations with adjuvant (either Alum or CpG) led to a higher mean mutation frequency in BM IgG (p<0.05 for OVA+CpG vs. PBS and p=0.07 for OVA+Alum vs. PBS).

Furthermore, Alum and CpG impacted the tissue-related mutation accumulation differently. There was a pre-existing tissue-related mutation frequency discrepancy, with BM consisting of sequences of a higher mean mutation frequency than that of spleen sequences independent of isotype (Fig. 2A and Fig.S4B). Upon immunizations in the IgG compartment, OVA alone decreased such a discrepancy by preferentially increasing mutations of splenic sequences (Fig S4B), CpG maintained it by increasing BM and spleen sequence mutations similarly (p<0.005, Fig S4B), and Alum enlarged it by accumulating more BM mutations (p<0.005, Fig S4B). On the other hand, in the IgM compartment, immunizations reduced the mutation frequency difference between tissues by significantly decreasing the mutations in the BM IgM compartment (Fig S4B). Such tissue-dependent immunization effects became more evident when we plotted the frequency discrepancy between BM and spleen across varying immunization conditions (Fig. 2D).

Overall, our data suggested: there were tissue- and isotype-related mutation frequency discrepancies at the steady-state, with higher mutations found in BM and IgG compartments; immunizations increased the mean mutation frequency of IgG sequences but decreased that of IgM sequences; OVA alone affected sequence mutations mainly in the spleen; the CpG immunization impacted mutations of BM and spleen sequences similarly; Alum more efficiently increased mutations of BM IgG sequences.

### IGHV gene usage and CDR3 length showed a tissue- and isotype-related difference and affected by adjuvanted immunizations

IGHV gene segment frequencies have been applied to distinguish stages of immune responses, describe age-related alterations, and define general B cell populations ^69, 70, 71^. In this work, instead of identifying altered frequencies of individual gene segments, we were interested in discovering correlated gene segment usage changes under different conditions. These changes could reflect a common or convergent development of Igs driven by biological or immunological regulators. The analysis started with identifying germline VDJ genes by a maximum likelihood algorithm developed previously by our group^38^. It is alignment-based and uses a gene-segment library derived from the IMGT^35^. Germline VDJ genes were annotated for each consensus sequence, and counts of different IGHV segments in each sample were obtained as the first step of the analysis. Then counts were transformed into compositions, an appropriate and required format for high-throughput sequencing data in microbiome studies^72, 73^. Next, we performed the principal component decomposition of the CLR transformed data. Principal component analysis (PCA) proved effective in capturing genuine biological differences among samples by relating genes with similar expression pattern^74^. We hypothesized that PCA of IGHV gene usage data produces a limited number of robust principal components, and many of them reflect the biological/immunological factor effects.

PCA on the IGHV gene usage data identified a group of six robust principal components (PC), each of which explained more than 5% of the total variations and altogether contributed ∼58% of the total (Fig. 3A). These PCs also had large Cronbach Alpha values (>0.75), which indicate a high internal consistency as a group. By relating IGHV genes of similar patterns, these PCs revealed underlying effects or regulations by factors such as immunization status, tissue location, or isotype identity (Fig 3B, C and Fig S7). PC1, explaining 15% of the total variations in IGHV gene compositions, reflected mainly an immunization effect (Fig 3D). The immunization effect was also tissue- and isotype-dependent (p=0.033 for the three-way interaction test in ANOVA). OVA alone affected PC1 mainly in the spleen IgG compartment, while OVA plus adjuvants (both CpG and Alum) resulted in significant changes of PC1 in the BM IgG compartment. Such a differential IGHV usage in different compartments upon adjuvanted immunizations indicated a possible mechanism underlying adjuvants’ effect, promoting homing of specific B cells to the BM niche. Furthermore, such an effect were most likely secondary during the immune response since the changes were limited to the IgG compartment. The top contributing IGHV segments to PC1 were highly correlated and almost exclusively contributed to PC1 (Fig. 3E and Fig. S8). They contained both up-regulated (IGHV2-2, IGHV5-4, IGHV5-12, etc.) and down-regulated gene segments (IGHV15-2, IGHV14-1, IGHV5-9, etc.) (Table S2).

**Figure 3.**
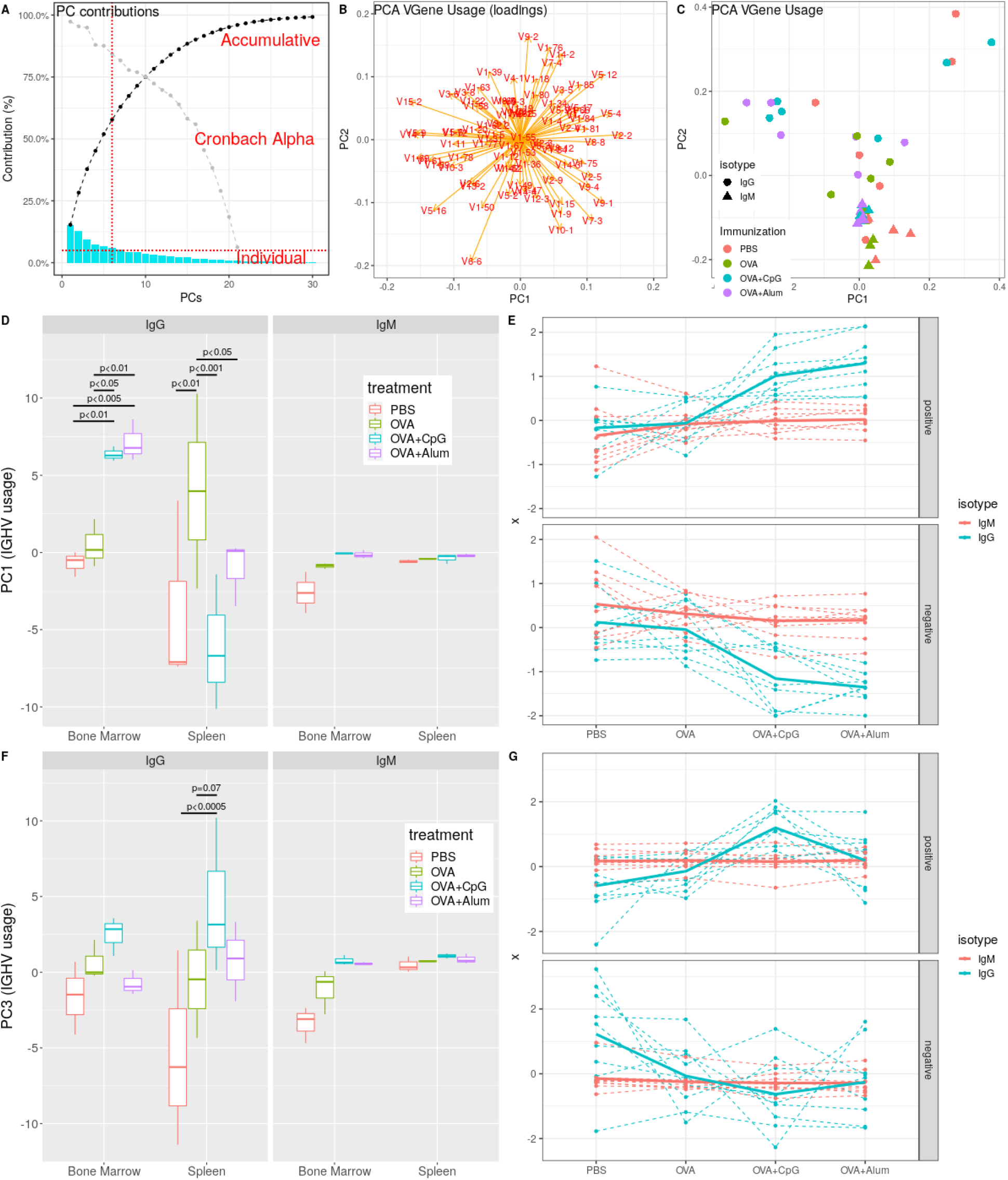
OVA immunization affects IGHV gene usage and adjuvants modify the tissue distribution of the effects. IGHV genes were annotated and gene usage was expressed as the percentage of each IGHV gene segment in each sample. The data were transformed into compositional data format. PCA was applied to reduce data dimensions and group correlating gene usages. Statistical analyses were done on top PCs with large contributions. A) The PCA scree plot showed the individual and accumulative PC contributions. The top six PCs with largest contributions (each explaining >5% total variation) have an accumulative contribution of about 58%. We also plotted the Cronbach Alpha value of each PC (grey dashed line), which is a measure of internal PC consistency. B) The PC loading plot. It showed correlations of IGHV gene usages projected on the top two PC plane. Genes are highly correlated if they point to the same directly, and contribute more to a PC components if they have larger loadings. C) Sample distribution on the first two PC space. The samples were labelled by their isotype (point shape) and immunization group (color). D) The PC1 distributions by different isotype, tissue and immunization status. PC1 explained about 15% of total variations and showed a significant immunization effect in a tissue- and isotype-dependent way. E) The raw IGHV gene usage patterns to verify the PC1 distribution. We plotted the top 10 raw gene usages (centered by the mean of each group) with largest contributions to PC1. Both positively (top) or negatively (bottom) contributing gene usages were shown. F) The PC3 distributions by different isotype, tissue and immunization status. PC3 explained about 9.5% of total variations and showed a significantly different pattern induced by the OVA+CpG immunization in the spleen IgG compartment. G) The raw IGHV gene usage patterns to verify the PC3 distributions. We plotted the top 10 raw gene usages (centered by the mean of each group) with largest contributions to PC3. Both positively (top) or negatively (bottom) contributing gene usages were shown.

Similarly, we found that PC2 (explaining 13% of the total variations) exhibited statistically significant isotype, tissue, and immunization-related changes. First, we found significantly different PC2 values between the IgG and IgM compartments (Fig S10A and C). The significant PC2 difference at the steady state indicated a naturally biased usage of this group of IGHV segments between different isotype compartments. Such isotype-related usage bias reached the maximum in the spleen at the steady state and became diminished upon immunizations or BM relocation (Fig S9C). In addition, the immunization effects on PC2 values were statistically significant only in the IgG compartment. The OVA+Alum immunization resulted in significantly increased PC2 values in BM IgG (p<0.001) but decreased PC2 values in spleen IgG (p<0.001, Fig S9C, D) compared with PBS controls and OVA alone samples. On the other hand, OVA+CpG only led to increased PC2 values compared with the PBS and OVA alone groups in the BM IgG compartment. Taken together, PC2 included a group of IGHV genes whose usages are normally biased between different isotypes and were impacted distinctly by the Alum or CpG immunization in different compartments.

Lastly, the PC3 group of IGHV genes (explaining 9.5% of total variations) demonstrated a different treatment effect induced only by the CpG immunization. In this case, OVA+CpG immunization resulted in a significant impact on IGHV usage in the spleen IgG compartment compared with other groups (Fig. 3F, G). Changes were also seen in the BM IgG compartment but without significance. The top contributing IGHV segments of PC3 consisted of up-regulated and down-regulated gene usages (Table S2).

CDR3, the most variable part of an Ig, is the principal determinant of the specificity and affinity of an antibody ^75^. Its length distributions and amino acid properties have been applied to characterize functional Ig repertoires, e.g., long CDR3s in the case of broadly neutralizing antibodies against HIV ^76, 77^. Here, we investigated how OVA immunization and adjuvants affected IGH CDR3 lengths. Interestingly, CDR3 length changes were similar to the IGHV gene segment usage results. First of all, statistically significant differences were limited to the IgG compartments upon immunization. OVA with either Alum or CpG resulted in significantly longer CDR3s than PBS or the OVA alone group in the BM IgG compartment (p<0.05, Fig 4), and OVA+CpG also led to an increased mean CDR3 length in the spleen IgG compartment (p<0.005, Fig. 4). As a result, under the OVA+Alum condition, we observed significantly longer IgG CDR3s in BM than those in the spleen. But in the case of OVA+CpG, we found equally long CDR3s in IgGs of both tissues. Lastly, CDR3 lengths in IgM compartments were not affected significantly by OVA immunizations. Again, these changes of mean CDR3 length were similar to the IGHV gene usage alterations but showed no correlations to the sequence mutation frequency (Fig. S13). One thing also worth mentioning was that at the steady state (PBS controls), CDR3s were shorter in IgG than in IgM compartments (p<0.05 for the IgG vs. IgM comparison in BM, Fig S12B), as were reported previously ^2^.

**Figure 4.**
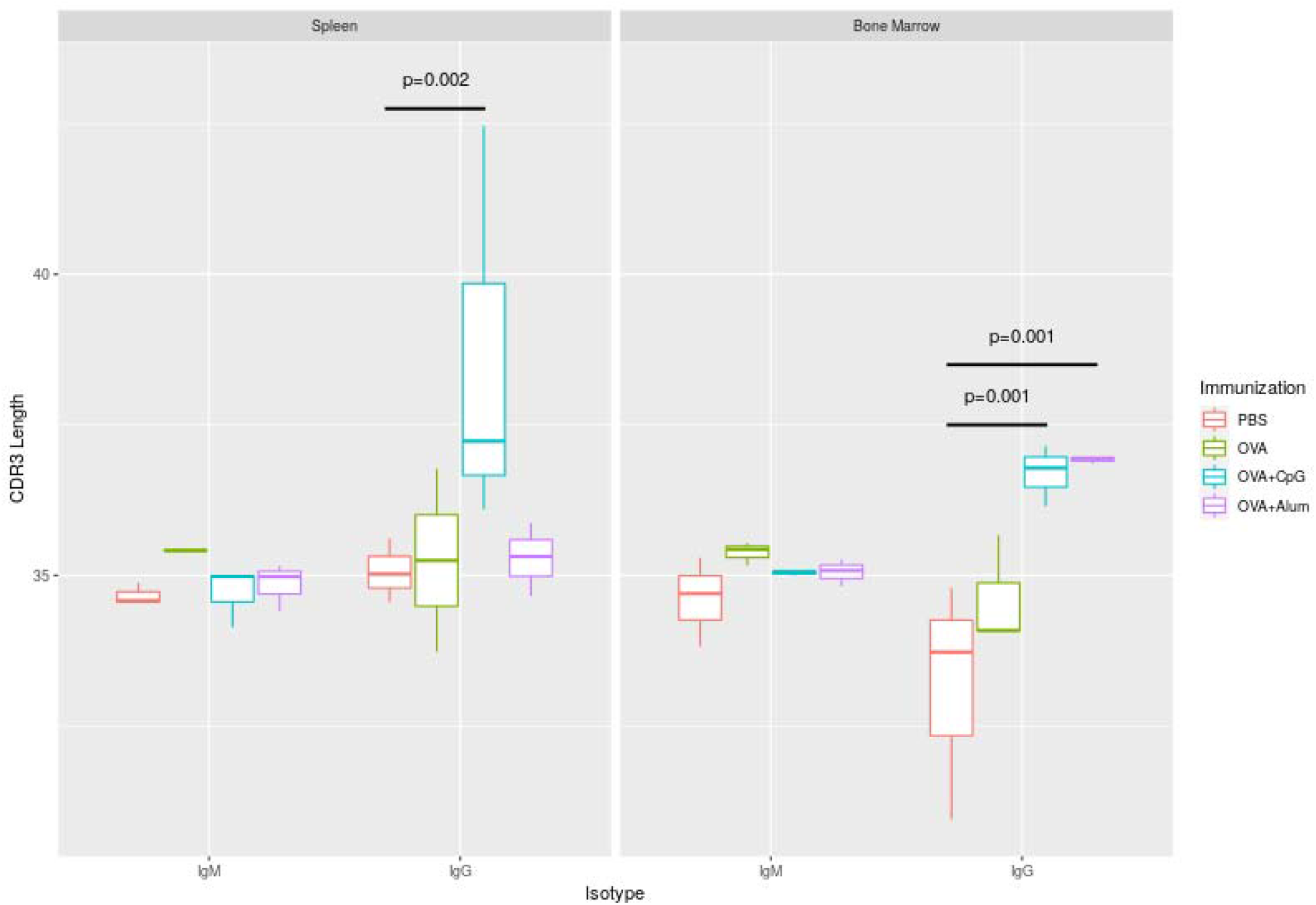
CDR3 length is affected by OVA adjuvant immunizations in a tissue- and isotype-dependent fashion. IGHV genes were annotated and CDR3 regions was identified. The CDR3 length distribution of each sample was determined, and the mean CDR3 lengths were compared and plotted by different compartments and immunization status. (Statistical p-values were corrected by FDR and significant p-values were indicated)

### Ig repertoires of different tissue-isotype compartments displayed distinct clone diversity profiles and CpG and Alum affected them differently

Diversity is a frequently used metric in ecology describing both the species variety and the relative abundance in an assemblage. Hill numbers, also known as the effective numbers of species, have been employed increasingly to quantify and assess biological diversity on the basis of sampling data^78^. They incorporate a mathematically unified group of diversity indices, differing among themselves only by an order ***q*** that defines the sensitivity of the number to the abundance or relative abundance of species (or clones in our work). When the order is small, diversity values are computed by weighting rare (small) clones more. As the order increases, diversities are calculated by weighting abundant (big or dominant) clones more^79^. Diversity of order zero, ^**0**^ ***D***, is insensitive to the clone size/abundance and referred to as the clone richness (i.e., total number of clones), ^**1**^ ***D*** is the well-known Shannon entropy, and ^**2**^ ***D*** is the Simpson concentration or the inverse Simpson index. In practice, ^**1**^ ***D*** is considered a measure of the effective number of common clones in a compartment, and ^**2**^ ***D*** or higher order diversities are of dominant clones ^40^. In this study, we worked on diversities with orders ***q*** between 0 and 4 since the diversity profile changes little beyond ***q*** =3 or 4^40^.

Clone abundance data, generated by the maximum likelihood estimation of clone lineages using Cloanalyst^38^, were used for estimating the asymptotic diversity Hill numbers. One concern about Hill numbers is that sample size affects the estimation: incomplete sampling underestimates the diversity values of low orders^40^. By resampling the clonal data at varying sample sizes, we showed that the diversity estimates are sensitive to sample size at low orders ***q*** <1.5 (Fig. S15). To overcome such bias, we determined the lowest sequence depth among all samples and then ran random subsampling 100 times of each sample with that number to estimate the diversity profile. With these profiles, we then started studying factor effects. First, upon comparing them between isotypes, we found that clone diversities of IgM compartments were much higher than those of IgG compartments at all orders, ***q*** =1 to 4 (Fig. 5A and Fig. S16A). However, when checking the clone size distributions, we found something contradictory. Most IgM compartments consisted of small clones with 1 or 2 members, but IgG compartments contained a significant proportion of larger clones (e.g., 15% of the clones in the spleen IgG compartment consist of more than five members) (Fig 5B and Fig. S14). Intuitively, we expected the IgG compartment to have more abundant clones, i.e., higher effective numbers or greater diversities at higher orders. The contradiction arose because diversity depends on two parameters: clone richness and clone evenness (relative abundance/size of clones); the diversity profile of the IgM compartment was more skewed by the high clone richness than it was in the IgG compartment. To control the effect of clone richness, we rescaled the diversity profile by dividing by its clonal richness (Fig 5C, D, and E). This way, we control for the effect of clone richness and make all diversity profile curves starting from the same point, ^**0**^ ***D*** = **1**. In addition, the rescaled diversity value at the order of 1 (^**1**^ ***D*** ) equals to the exponential of the Pielou evenness index.

**Figure 5.**
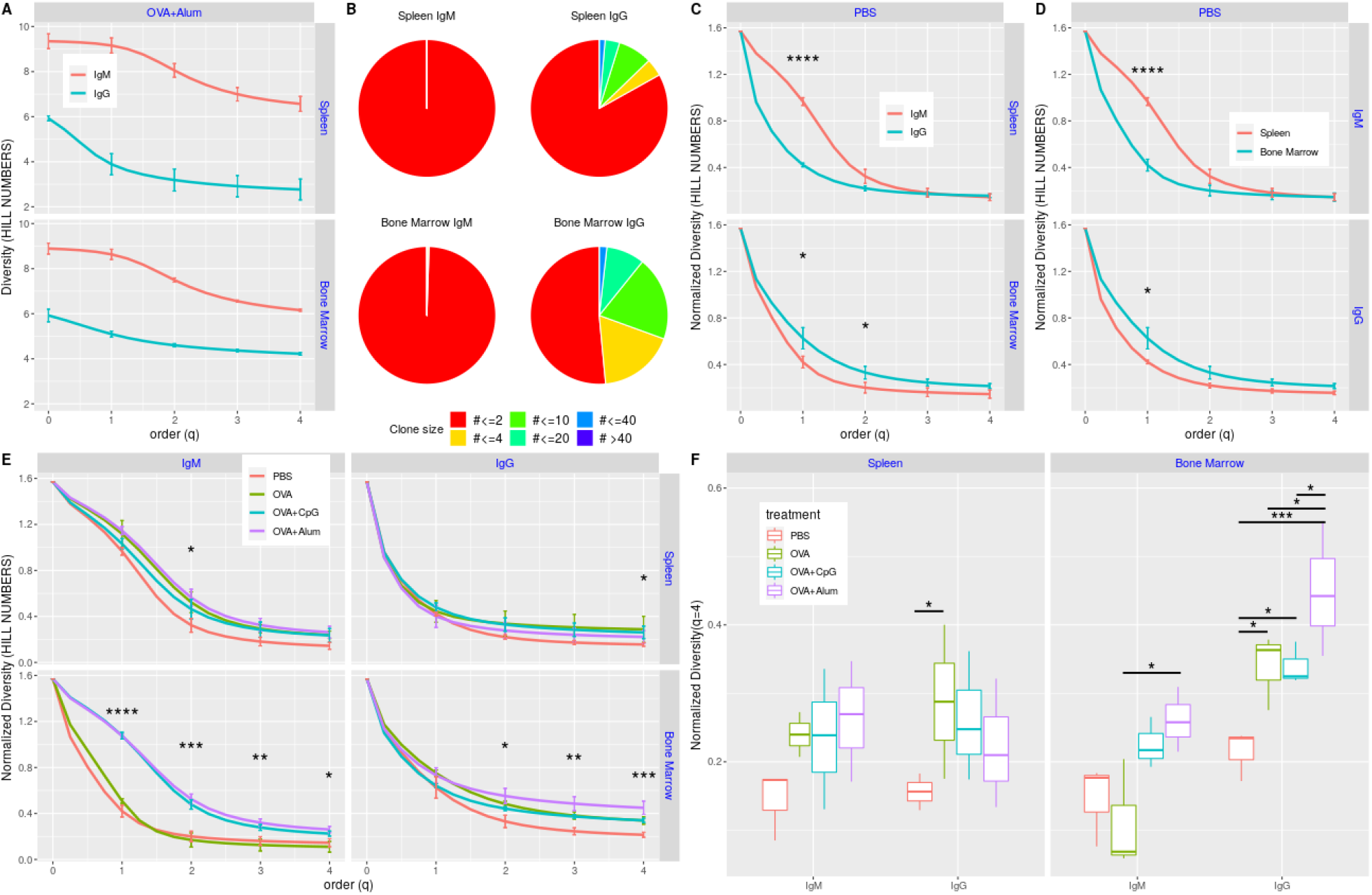
Clonal diversity as Hill numbers. Ig sequence data were annotated and partitioned into clones. Diversity profiles as Hill numbers were estimated based on clone abundance data with orders between 0 and 4. They were also scaled to control the effect of clone richness and focus on clone evenness (relative clone abundance/size). A) Diversity profiles of different isotype compartments under the OVA+Alum immunzation. Each line is for one isotype compartment in different tissue. Standard errors are plotted for diversities at the order of 0, 1, 2, 3 and 4. B) Example clone size distribution pie charts. Here shows the data for one mouse immunized with OVA+Alum, and results for four compartments of different isotype and tissue combinations were shown. C) Scaled diversity profiles of different isotype compartment at the steady state. Each line is for one isotype compartment in different tissues. Standard errors are plotted for diversities at the order of 0, 1, 2, 3 and 4. D) Scaled diversity profiles of different tissue samples at the steady state. Each line is for one tissue sample in different isotype compartments. Standard errors are plotted for diversities at the order of 0, 1, 2, 3 and 4. E) Scaled diversity profiles of samples under different immunization status. Each line is for one samples under different immunization in different tissue-isotype compartments. Standard errors are plotted for diversities at the order of 0, 1, 2, 3 and 4. F) Scaled diversity distribution and comparison of diversity values at the order q=4 for samples under different immunization conditions in different tissue-isotype compartments. (Statistical p-values were corrected by FDR and significant p-values were indicated. *, p<0.05; **, p<0.01, ***, p<0.001, ****, p<0.0001)

Clone diversity profiles before or after rescaling exhibited quite distinct patterns when comparing repertoires of different isotypes, tissues, and immunizations (Fig. S17 and Fig. 5E). We focused on the rescaled diversity profiles to be less confounded with clone richness. We first compared the steady-state (PBS controls) diversity profiles between different isotypes and tissues. In spleen, higher diversity values were observed for IgM clones than those of IgG clones; in BM, it was IgG clones showing higher diversities. When comparing between different tissues, BM clones exhibited higher diversities in the IgG compartment, and spleen clones were of higher diversities in the IgM compartment (Fig. 5C, D). Furthermore, the observed difference was statistically significant only at the small order q=1 or 2, indicating changes occurred to clones of smaller size.

Next, OVA alone increased clone diversities compared to the non-immune state profiles, with a tissue and isotype dependency. For IgM clones, OVA alone increased diversities only in the spleen but only significantly at smaller orders (p<0.05 at q=1 or 2), while for IgG clones, it increased diversities significantly at larger orders and in both spleen and BM (p<0.05 at q=4 in spleen and at q=2, 3, 4 in BM, Fig. 5E, F and Fig. S17). The OVA alone immunization did not change the pattern of diversity difference between tissues or isotypes observed at the unimmunized state but simply resulted in enlarged gaps (Fig. S17A). Under OVA alone immunization and compared between isotypes, the IgM diversity profile showed significantly higher values (p<0.001) for small orders but slightly lower values than the IgG profile for large orders in the spleen (Fig. S17A); the IgG profile was significantly higher at all orders (P>0.05 for all comparisons at q=1 to 4) in BM (Fig. S17A). Additionally, when compared between tissues, the spleen clones showed significantly higher diversity values in the IgM compartment than the BM clones for all orders (p>0.05 for all comparisons); it was the BM that contained significantly higher diversity values in the IgG compartment than the spleen but only for smaller orders q=1 (p<0.05, Fig. S17B).

Similar to OVA alone, in the spleen, OVA+Alum increased diversities of IgM clones at small orders. However, it affected spleen IgG diversities to a lesser degree (Fig. 5E and Fig. S17). Significant changes induced by OVA+Alum came from BM clone diversity profile. It resulted in significantly higher BM clone diversities – for both the IgG and IgM isotype – than those of OVA alone or PBS groups (Fig 5E, F and Fig. S17). As seen above, the increases in IgM clone diversities were mainly at smaller orders (p<0.0001 for all comparisons at q=1 and 2), and the increases in IgG were at higher orders (p<0.05 at q=3 and 4). Consequently, the IgM clone diversity profiles in spleen and BM become almost identical when compared between tissues (Fig. S17B), while BM had higher IgM clone diversities at smaller orders but higher IgG clone diversities at higher orders when compared between isotypes (Fig. S17A).

Finally, the OVA+CpG immunization gave rise to diversity profiles similar to those yielded either by OVA alone or by OVA+Alum, depending on the tissue and isotype. It raised BM IgM diversities significantly at lower orders resulting in a similar profile to the one caused by OVA+Alum, while for the IgG diversities, it led to profiles more resembling the ones by OVA alone with increasing values at higher orders in both BM and spleen (Fig. 5C, D, E, F and Fig. S17).

### Assessing clone selection strength and adjuvants’ effects on it

We developed a method to assess the level of selection a B cell clone experienced throughout its development. It involves comparing two parameter estimates, mean mutation frequency and intra-clonal pairwise sequence dissimilarity. The rationale is that a clone evolving under selection expands some members with a high affinity for the specific driving antigen; thus, it grows more mutations but less dissimilarity among those. Otherwise, it evolves with both parameters advancing proportionally. This reasoning can be verified by visualizing two parameters on an x-y plot (Fig. S19). Under a naïve condition (the IgM compartment in PBS control), the two parameters of most clones showed values in good proportion, and clones settled on the plot following a straight trend line (Fig. S19). Based on such a distribution, we set up a threshold to separate clones. Clones in the first group stayed above the threshold line and had a high mean mutations frequency and high sequence similarity, representing clones likely being selected through its development. In the second group, clones were located below the threshold and consisted of members with balanced mutations and dissimilarity, representing clones evolving under no selection. To quantify and compare the clone selection metric among samples, we selected the top 40 clones (based on the size)^80^ from every sample and calculated the percentage of clones under selection (above the defined threshold on the two-parameter plot) (Fig. S19 and Fig. 6A, B). Then we carried out statistical analyses on these percentage data.

**Figure 6.**
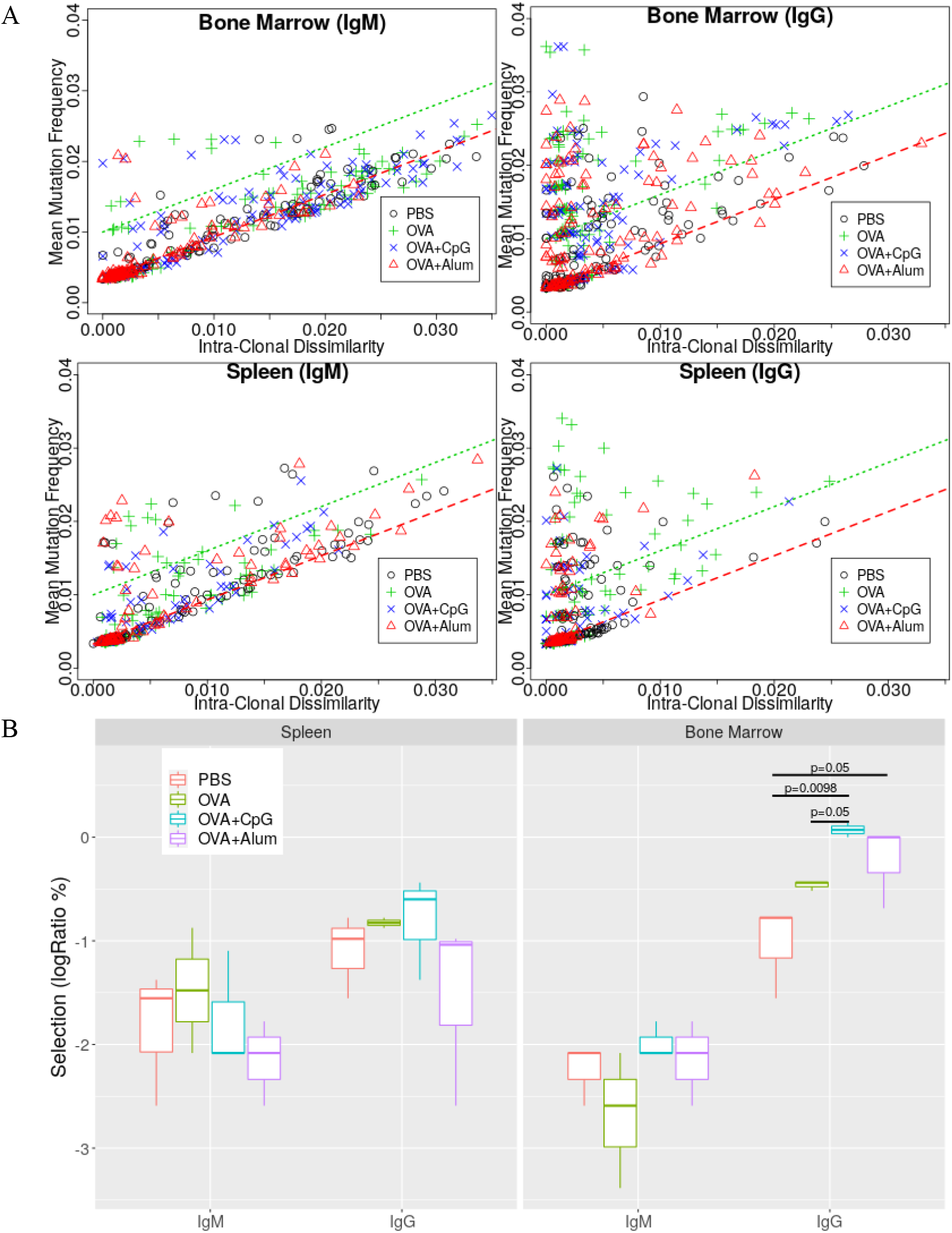
Evaluation of clonal selections and their changes by OVA immunization with adjuvants. We developed a new metric to quantify the changes of clone phylogenetic trees to reflect the effect of selection. It involves two measurements, mean clonal mutation frequency and pair-wise intra-clonal dissimilarity. Under no selection, a clone evolves to increase these two parameters in proportion, and two parameters show as a straight line on an x-y 2D plot. Otherwise, some parts of a clone tree is selected, and the linear relationship between the two parameter is distorted. To analyze and compare the selection strength metric, we plotted the two parameters of largest 40 clones of each sample (A) and defined a threshold (the green dashed line), above which are clones showing some level of selection. For each sample, on the basis of such threshold we can obtained the percentage of selected clones among the largest clones. The percentages were compared statistically and results were shown in (B). (Significant p-values are showed for comparing OVA immunization effects)

Under all immunization conditions and in both tissues, the IgG compartment included a significantly higher percentage of clones under selection than the IgM compartment (Fig. S20A). Such observation matched our expectation since IgG B cells were memory cells educated through the germinal center reaction; IgM B cells were either inexperienced naïve cells or exited early from germinal centers during an immune response ^65, 81^. Furthermore, the BM IgG compartment contained a significantly higher percentage of selected clones than the spleen IgG compartment either in PBS controls or under immunizations (Fig. S20B). These clones in BM most likely consisted of PCs exited from the germinal center at the later stage of an immune response and migrated to the BM niche for long-term survival and antibody production^65, 82^. Lastly, adjuvanted immunization (Alum or CpG) significantly increased the percentage of clones comprised of selected members in BM IgG compartments, with CpG resulting in the highest proportion (p<0.01 and p<0.05 for OVA+CpG compared with PBS and OVA alone, respectively, and p<0.05 for OVA+Alum vs. PBS) (Fig. 6B).

### Unsupervised machine learning to characterize Ig repertoire profile of different isotypes, tissues, and immunizations

We used unsupervised machine learning to identify associations among the various features of the Ig repertoire and the immunizations. The data are high-dimensional and of varied type: each sample is characterized by isotype, tissue of origin, and immunization status, as well as by mean mutation frequency, usage frequency over more than 130 V-gene segments, mean CDR3 length, Hill numbers of order 0-4, and mean selection strength. To deal effectively with such heterogeneous data, we first carried out a multiple factor analysis (MFA), a PCA-based multivariate data analysis intended to balance the influence of different variable types (quantitative or qualitative) by groups^83^. Without this balancing, the analyses would, for example, be dominated by gene segment usage.

The MFA transformed these variables into 15 principal components, the top two of which accounted for about 60% of the total variation (39% and 20% for PC1 and PC2, respectively; Fig. S21). We used unsupervised clustering on the data in transformed PCA coordinates. Multiple clustering algorithms on all 15 PCs were carried out, including agglomerative hierarchical clustering (HCL), partition around medoids (PAM), spectral clustering, and affinity propagation clustering (APC). All algorithms yielded similar results (Figs. S22 and S23).

Projection onto PC1 and PC2 shows that IgG+ and IgM+ samples are well-separated across PC1, while spleen and bone marrow divide roughly along PC2 (Fig. 7A and D). In the IgG+ compartment, the samples are separated by tissue specificity, but such separation is not evident among the IgM+ samples (Figs. 7A and7B). The impact of different immunizations depends strongly on tissue and isotype (Fig. 7C). In the BM IgG compartment, in particular, samples of different immunization groups aggregated into distinct clusters separated from each other (Fig. 7C and Figs. S22 and S23). In addition, the samples appear to be roughly ordered across PC1, starting with the PBS samples at the left, then moving right, OVA, and then the two adjuvanted immunization groups together. In the BM IgM compartment, we mainly saw two clumps, one for samples immunized in the presence of adjuvants (CpG and Alum) and the other for PBS and OVA alone. Patterns in spleen compartments were less clearly marked, with seemingly OVA alone and OVA+CpG groups organized into separated clusters (Fig. 7C and Figs. S22 and S23)

**Figure 7.**
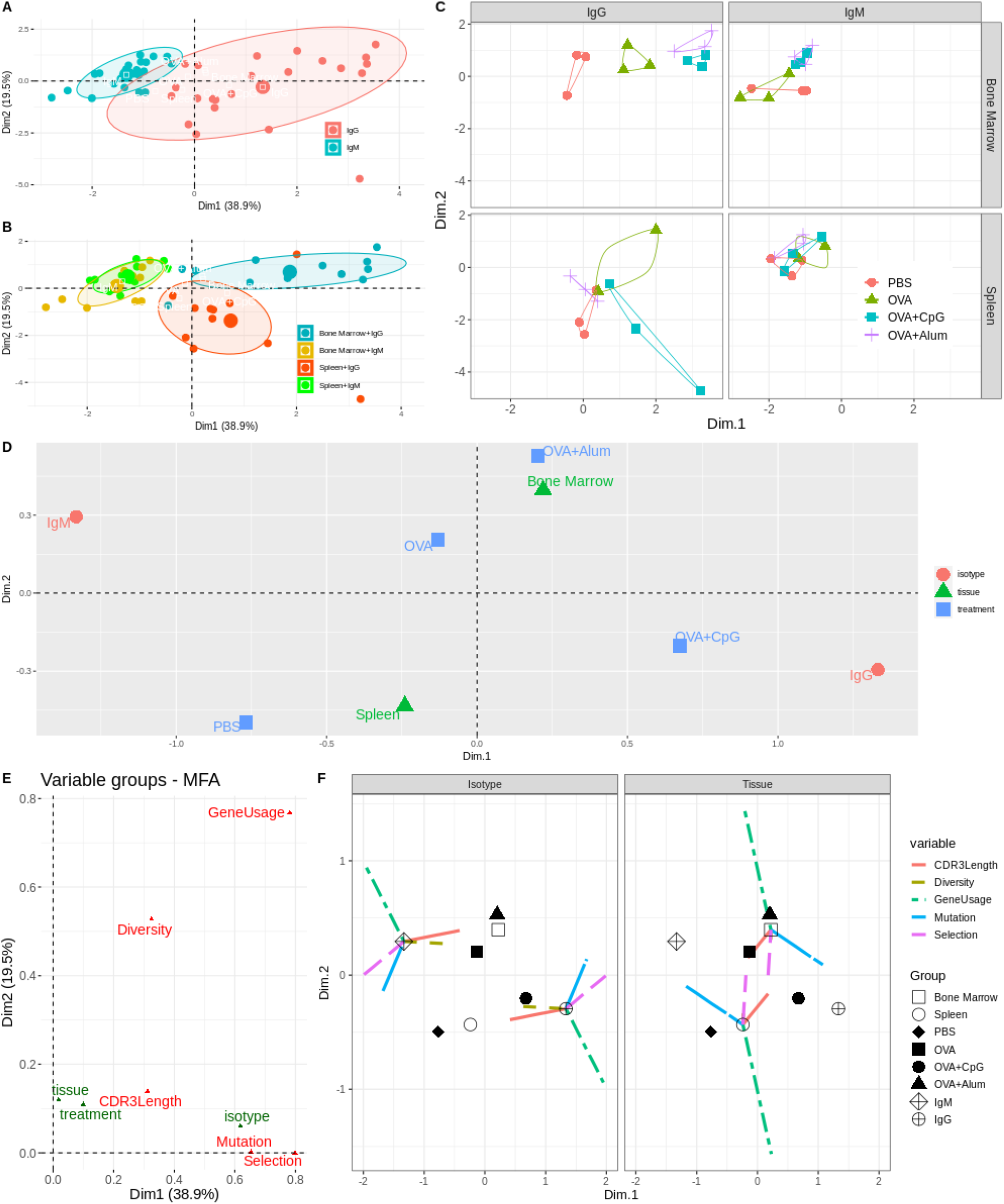
Unsupervised machine learning to characterize Ig repertoire changes. Multiple factor analysis (MFA) was carried out on the groups of variables including IGHV gene usage, mutation frequency, CDR3 length, clone diversity (expansion) and clone selection level. Sample data were then projected on the first two dimensions of MFA arranged by their isotype (A), tissue location (B), and immunization conditions (C). MFA can also summarize and project supplementary category variables (isotype, tissue and immunization) into the new PC dimensions (D) to show their correlations. Furthermore, the correlations between the variable groups and the first PC dimensions were shown in (E). Finally, the partial point graphs of supplementary category variables in MFA were shown in (F), in which supplementary categorical variables (such as isotype, tissue and immunization) were plotted as points with different shapes at the barycenter and variable groups were connected by lines with each center. The figure was arranged into sub-plots by isotype, tissue (F) and immunization (Fig. S26) for easy visualization.

Next, we projected supplementary categorical variables (i.e., grouping variables such as isotype, tissue, and immunization) on new PC dimensions to compare average changes among different categories. MFA allows for such projections by averaging information across samples in each category Fig. 7D shows the projections of these category variables on the top two PC plane. First, same as seen above, the isotype variable showed the largest variation, i.e., IgG and IgM categories stayed far separated with much distance, primarily along the PC1 direction. The tissue variable was distributed more along the PC2 dimension, almost orthogonal to the isotype variable (the PC1 dimension). The immunization variable correlated with tissue along the PC2 direction, except for the OVA+CpG category. The OVA+Alum group stayed very close to BM but equally distant to IgM and IgG, meaning the changes induced by OVA+Alum were more in BM and almost equally of both isotypes. On the other hand, the OVA+CpG group resided very close to IgG and slightly more adjacent to BM, meaning the changes caused by OVA+CpG were more of the IgG isotype and equally in both tissues. Such a pattern of changes matched what we saw in the individual sample projection data (Fig. 7A, B, and C). The OVA alone group stayed more adjacent to BM and IgM, meaning the changes of this category occurred more in BM and IgM compartments. However, it was also close to the origin of the coordinates, indicating its effects were lower in magnitude than other groups. Lastly, the PBS category remained more adjacent to IgM and spleen, meaning without vaccination the Ig repertoire contained alterations mainly in the spleen IgM compartment. Furthermore, we extended our analyses and observations to the top 3-PC space (altogether explaining about 78% of the total variations, Fig. S21) and found a similar pattern of changes (Fig. S24 and S25).

Isotype, tissue, and treatment associations with the natural variables was investigated by projecting each variable group onto specified planes in the PC space (Fig. 7E). This procedure revealed that PC1 is dependent largely on gene usage, mutation, and selection and PC2 depends on gene usage and diversity. In this way, we found that isotype, separated in PC1, was correlated with gene usage, mutation, and selection. Tissue of origin, separated in PC2, was correlated with gene usage and diversity (Fig. 7E and Fig. S26)). Finer detail can be found by Partial Point Analysis ^84^ (Fig. 7F and Fig S27). For example, although the clone selection levels of the BM IgG and BM IgM compartments were quite different from those of the respective spleen IgG and spleen IgM compartments (Fig. 6B), the overall clonal selection levels of BM and spleen were similar after averaging those of different isotypes in each tissue. That emphasized the necessity of studying Ig repertoires in separated tissue-isotype compartments.

Similarly, when analyzing treatment groups, we find that the PBS control and OVA immunized groups (with or without adjuvants) varied in diversity, CDR3 length, and gene usage, but not in mutation frequency. The apparent similarity of the clone selection levels between the PBS group and the other categories resulted from averaging over isotype. Differences between OVA alone and adjuvanted OVA immunizations also were attributable mainly to diversity and gene usage changes. When comparing OVA+CpG with OVA+Alum, all variable groups contributed similarly with almost equally distant partial points, except for the mutation frequency variable (Fig. S27), indicating they differed in IGHV usage, CDR3 length, diversity, and clone selection levels but exhibited similar mutation frequencies.

## Discussion

We developed an in-house Ig sequencing library preparation and data analysis pipeline for accurately profiling immunoglobulin heavy chain repertoires. The library preparation protocol is simple and generates UMI-labeled isotype-aware reads of high quality and, along with the subsequent data processing, increases the yield of unbiased Ig sequences. One important way in which this yield is enhanced is, in contrast with the common practice^9, 56, 57^, to omit paired-end joining. This increased the yield of usable reads 10-fold and eliminated errors caused by spurious short overlaps during paired-end joining. In addition, we developed new methods and metrics and adapted others to characterize the Ig repertoire: simplifying IGHV gene usage information using PCA, introducing diversity profiles using rescaled Hill numbers, and developing a two-parameter clonal selection metric based on the intra-clonal distribution of mutations. We combined these analyses with unsupervised machine learning to characterize the effect of immunization in different tissue-isotype compartments and the contribution of repertoire properties to that effect.

We applied these techniques to characterize the impact of two commonly used adjuvants, CpG and Alum, on the murine heavy chain Ig repertoire in an OVA immunization model. We collected data from four compartments distinguished by tissue (BM or spleen) and isotype (IgM or IgG). The hypothesis was that these compartments contain functionally and phenotypically distinct B cells, and that the different immunization formulations affect them differently. Indeed, we found that the Ig repertoires from different tissue-isotype compartments differed in many respects and that immunization had a further tissue- and isotype-dependent impact. Isotype accounted for the highest variation in the repertoire, with 40% of the overall variance. IgM and IgG repertoires differed in IGHV mutation frequency, selection level, and IGHV gene usage. At the steady state in PBS controls, the IgG repertoire consisted of sequences using different IGHV segments and showing higher mutation frequency and contained more clones experiencing selection than the IgM repertoire. Tissue and immunization jointly resulted in changes explaining about 20% of the overall variations and orthogonal to the isotype effect. Repertoires from different tissues differed in IGHV gene usage and mutation frequency. At the non-immune steady state (PBS controls), the repertoire from BM consisted of sequences utilizing different sets of IGHV genes and accumulating more mutations than the spleen repertoire. These repertoire changes related to isotypes or tissues matched what we expected based on functional and phenotypic properties of B cells in these compartments18, 65, 66, 67, 68.

Interestingly, distinct immunization conditions altered the Ig repertoire differently, and their effects exhibited different tissue and isotype dependencies. The non-immune naïve repertoire (PBS controls) showed pre-existing tissue and isotype bias in gene usage, mutation frequency, and diversity, as mentioned above. Moreover, compared to OVA immunized repertoires (either adjuvanted or OVA alone), the naive repertoire utilized a distinct group of IGHV gene segments and comprised more rare clones (higher diversity values of small orders). The OVA alone immunization resulted in a repertoire different from the naive one, and its effect was more pronounced in BM or the IgM compartment. Repertoires under two adjuvanted immunizations differed from those under PBS and OVA alone immunization and distinguished from each other as well. OVA+Alum led to a repertoire profile staying near BM and in roughly an equal distance to both isotypes on the MFA factor map, indicating it causes changes more in BM of both IgG and IgM compartments. However, OVA+CpG resulted in a profile close to the IgG isotype and equally distant to both tissues on the factor map, meaning it invoked alterations in the IgG compartment of both tissues. Overall, OVA alone could accumulate significantly more mutations than the PBS control, to a level slightly lower than adjuvanted immunization did. Compared with OVA alone, adjuvanted immunizations did not impact the gene mutation but changed gene usage, clone diversity, and clone selection of the Ig repertoire significantly. Alum resulted in more abundant clones in BM for both isotypes, presumably by promoting the BM migration of these clones or survival of them in BM. On the other hand, CpG more efficiently increased the clone selection but affected to a lesser degree the diversity of abundant clones and exerted its effect on the IgG compartments in both spleen and BM. Based on these observations, we proposed distinct adjuvant effect models (the quantity vs. quality model, Fig. S28). These two models are not mutually exclusive and need more evident to confirm.

In summary, we characterized the Ig repertoire profiles in different compartments and under varying immunization conditions. These observations were not surprising but confirmed previous functional and phenotypic studies. In addition, we found that CpG and Alum resulted in distinct Ig repertoire profiles, probably due to differences on the cell types they affect and their mechanisms of action (via TLR9 for CpG vs more of a non-specific effect for Alum). Looking ahead, we will want to expand the list of properties of Ig repertoire to be studied, such as CDR3 amino acid composition, IGHJ gene usage, etc., along with comparison to both TLR dependent and independent adjuvants. In addition, there are more B cell compartments contributing to protective immunity depending on the route and location of vaccine or infection, such as draining lymph nodes, mucosal B cell, and gut and other tissue-resident B cells. Future work should characterize those repertoires, ideally at different time points during an immune response and including other isotypes. The pipeline we have developed can be easily applied to study the Ig repertoire of other species (human, rat, and so on).

## Abbreviations

Ig: immunoglobulin
IGH: immunoglobulin heavy chain
IGHV: heavy chain variable gene segment
IGHD: heavy chain diversity gene segment
IGHJ: heavy chain joining gene segment

## Notes

### Competing Interest Statement

The authors have declared no competing interest.

