## Supplementary Material File 1 for "Characterizing adjuvants’ effects at the murine immunoglobulin repertoire level"

Supplementary Materials

Library Prep

Figure S1 Quality

| A: Read 1 |  |
| --- | --- |
| 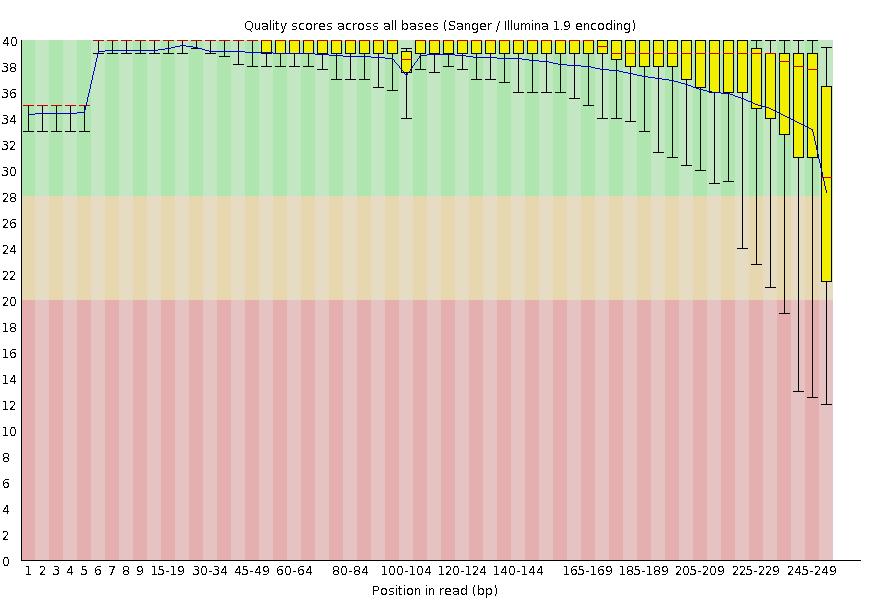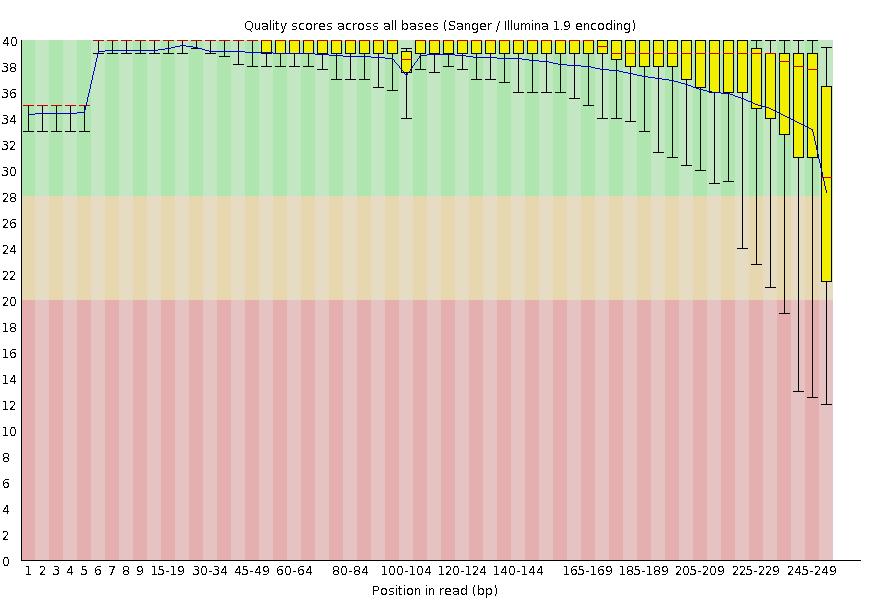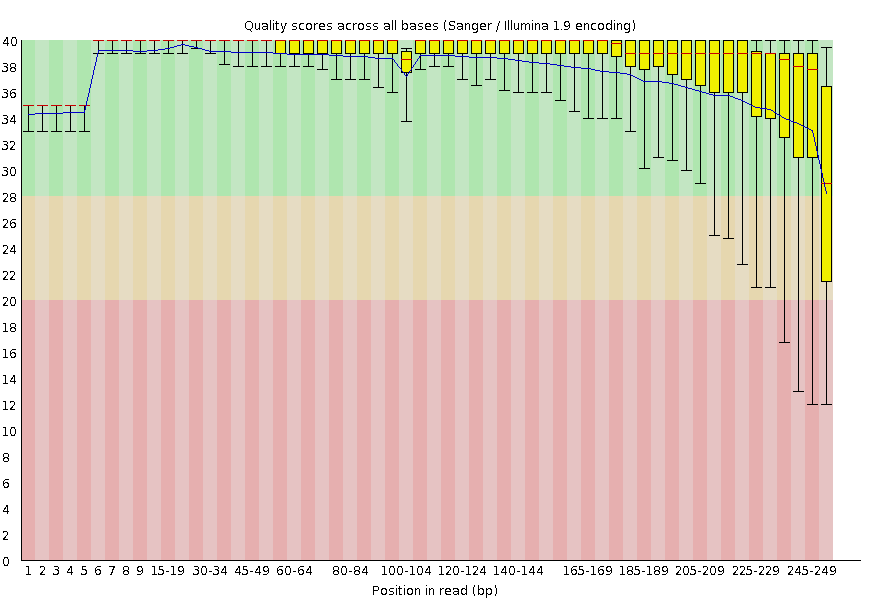B: Read 2 | 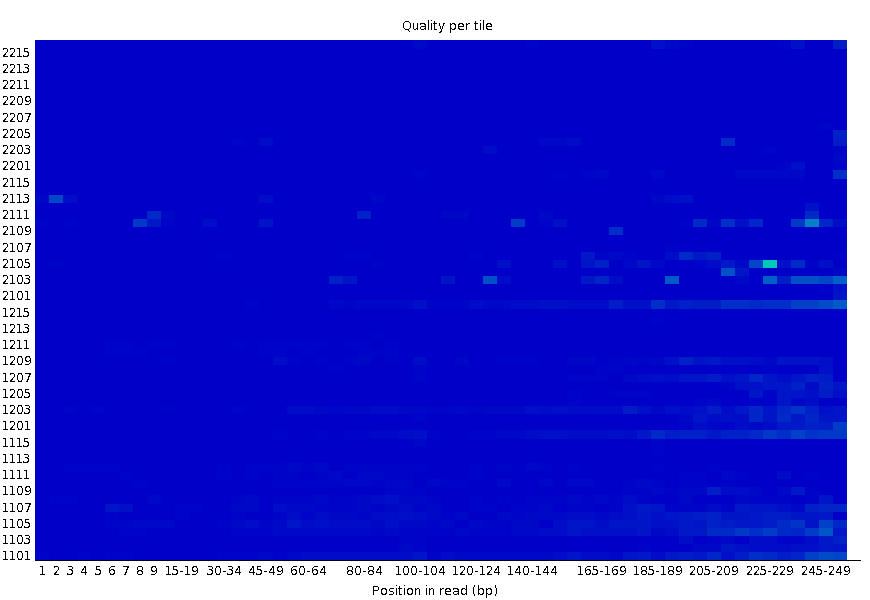 |
| 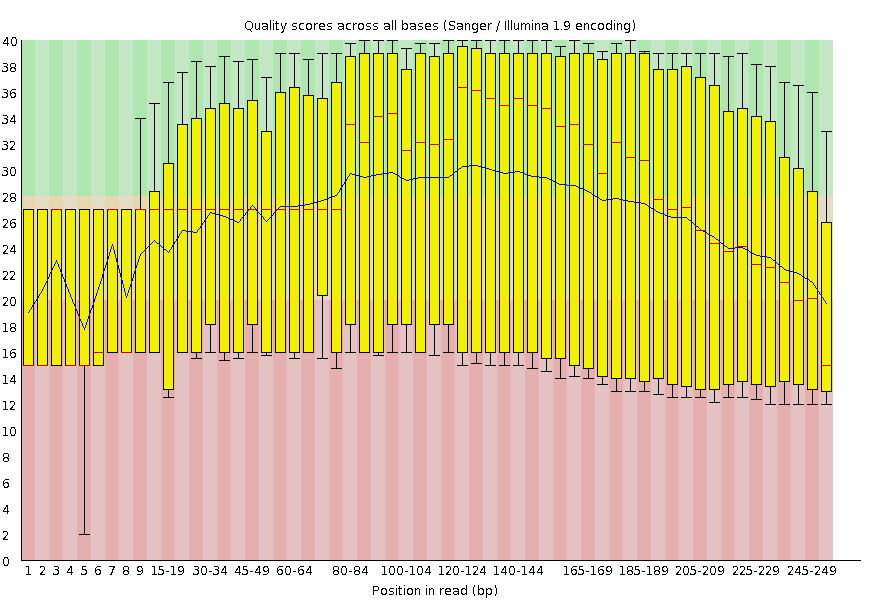 | 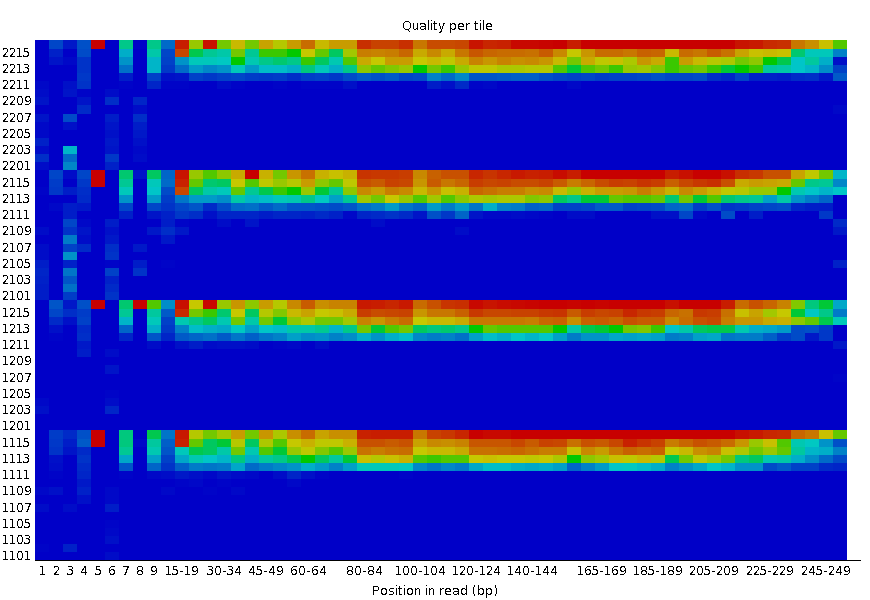 |
| C: Read 1 |  |
| 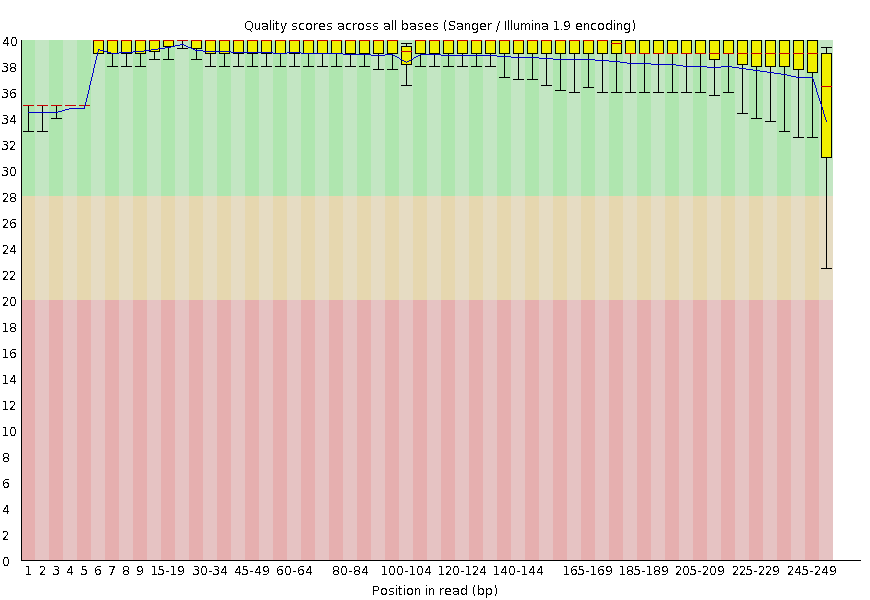 | 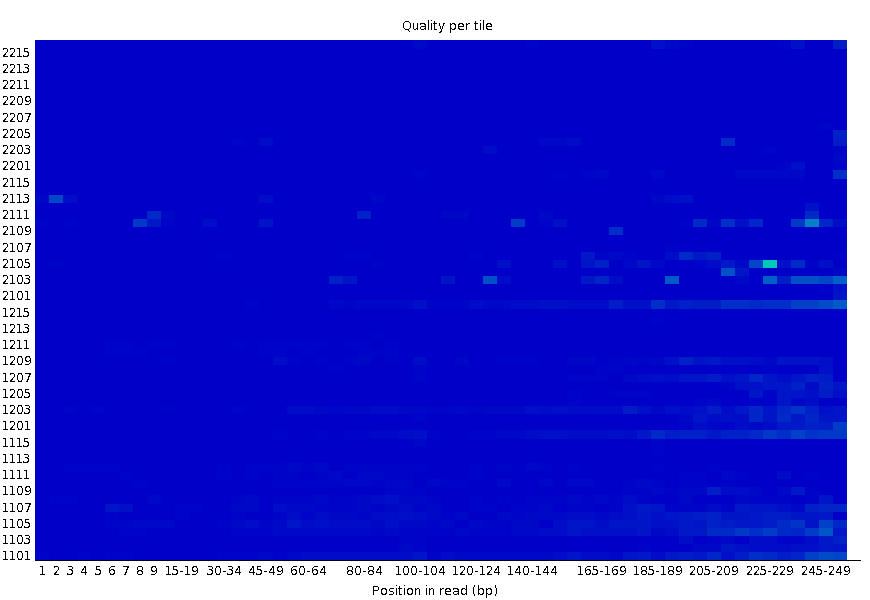 |
| (Figure S1 continued)  D: Read 2 |  |
| 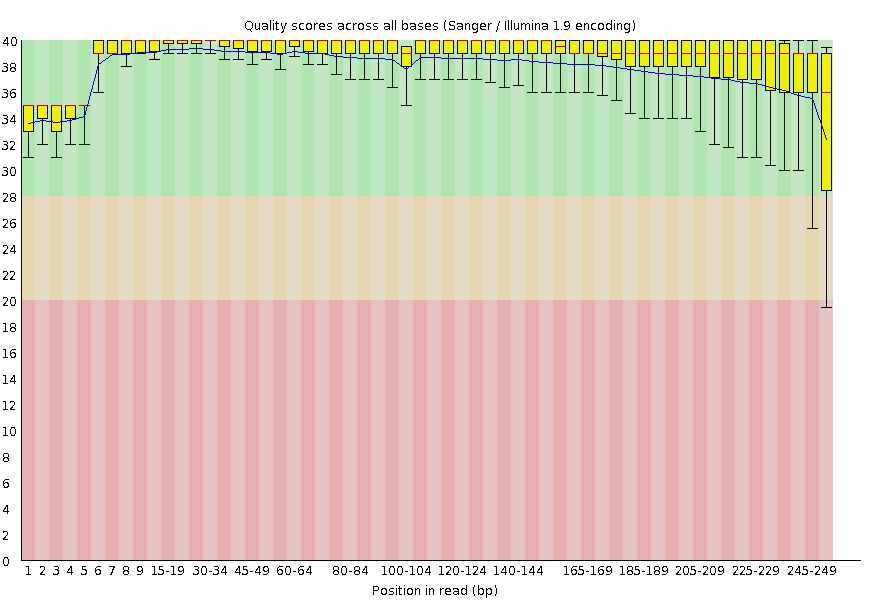 | 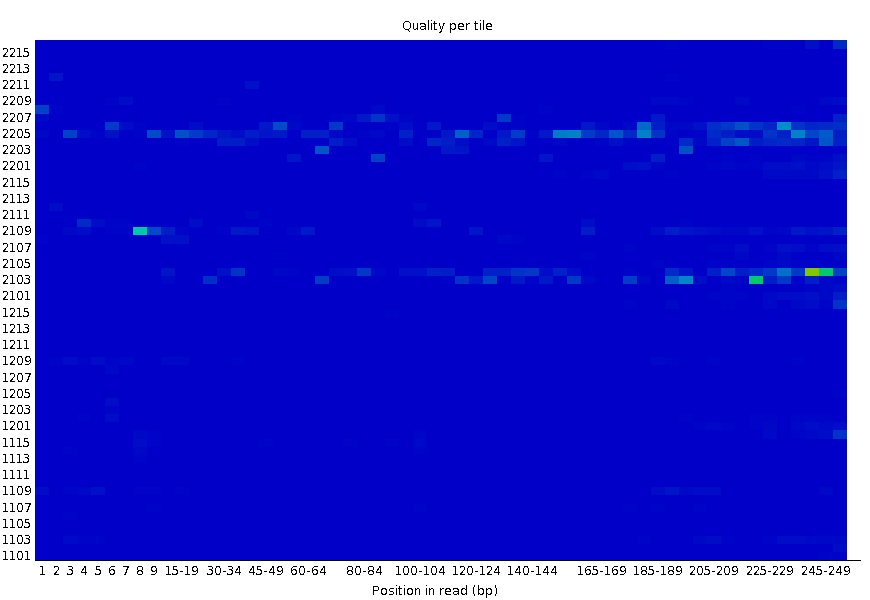 |

Figure S1. Example sequence QC plots. The plots were generated using FastQC (<https://www.bioinformatics.babraham.ac.uk/projects/fastqc/>). We showed the example per base sequence quality plot and per tile sequence quality plot for paired reads generated by the two protocol with IG constant region primers with or without random nucleotides. (A) and (B) QC plots for paired reads generated using the IG constant region primers with random nucleotides at the 5’ beginning end. (C) and (D) QC plots for paired reads generated using the IG constant region primer without random nucleotides at the 5’ end. ((A) and (C) are read 1 sequences; (B) and (D) are read 2 sequence). The IG constant region primers with random nucleotides helps to increase the diversity of sequences at the 3’ end and greatly improve the read 2 sequence quality, since the constant region and J chain of immunoglobulin heavy chain genes contain less diverse sequences.

Table S1. Summary statistics for the sequencing run of libraries prepared with isotype primers with or without 5’-random nucleotides. The quality and yield of the sequencing run with random nucleotides are much higher than the one prepared with the regular isotype primer.

|  | **Isotype-specific primer** | **Isotype-specific primer with 5’- random nucleotides** |
| --- | --- | --- |
| % PF Clusters | 98.11% | 99.09% |
| % >= Q30 Bases | 68.59% | 96.72% |
| Mean Quality Score | 32.28 | 38.57 |
| PF Clusters | 422,138 | 1,450,749 |

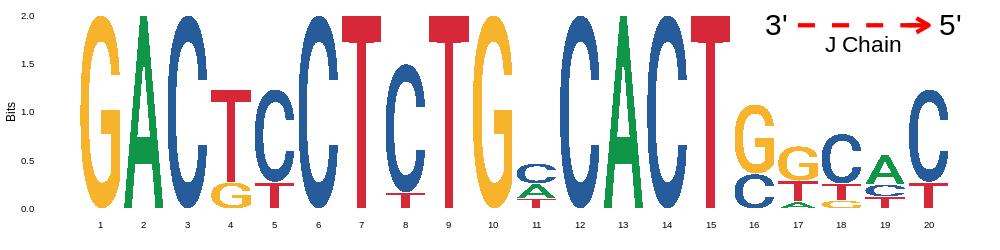

Figure S2. Conserved mouse IGHJ chain sequences. First twenty nucleotides of mouse IGHJ chain (3’→5’) were plotted as a sequence logo chart. It shows conservancy at multiple locations.

**UMI de-duplication and consensus reads generation**

Unique molecular identifiers (UMI) are widely used in sequencing to remove PCR amplification bias. There are many tools/algorithms to de-duplicate reads on the basis of alignment coordinates and UMIs in regular RNA sequencing work (Ref: <https://bmcbioinformatics.biomedcentral.com/articles/10.1186/s12859-019-3280-9>; https://www.ncbi.nlm.nih.gov/pmc/articles/PMC6921982/). However, not many tools has been specialized to process IgSeq data. Different from regular genes, Ig genes show a much higher level of sequence similarity for they are generated by recombination from sets of limited gene segments. Furthermore, sequences of high somatic hyper-mutations make the determination of alignment coordinates and error-correction sometimes impossible. Therefore, we developed a new deduplication pipeline that is tailored to work on IgSeq reads for collapsing duplicates and removing amplification bias and PCR/sequencing errors as well.

Our deduplication pipe include the following steps:

1) Group reads on the basis of UMIs and first 50 nucleotides at the 5’ end of sequences;

2) Refine UMI groups by accounting for sequencing errors in UMI sequences using the network-based algorithm^1^;

3) Align multiple sequences in each UMI group;

4) Refine UMI groups by checking constant region sequences at the 3’-end and separate the sequences on the basis of isotypes;

5) Split UMI groups on the basis of multiple alignment results allowing less than a 5% mismatch rate of each group and the mismatch rate is determined by a sliding window of 30nts;

6) Generate one consensus read for each UMI group in probabilistic way with the consideration of quality scores of each nucleotide;

Also in the pipeline, we allow users to remove singleton UMI groups (a UMI group containing only one sequence) and only generate consensus reads for UMI groups with at least two sequences.

The front end of the program implementing the new pipeline is implemented in python and it depends on few other packages in python, C++ and Linux utility to perform the jobs and tasks related to above steps. The code are available at the Github repository <https://github.com/BULQI/IgSeq>.

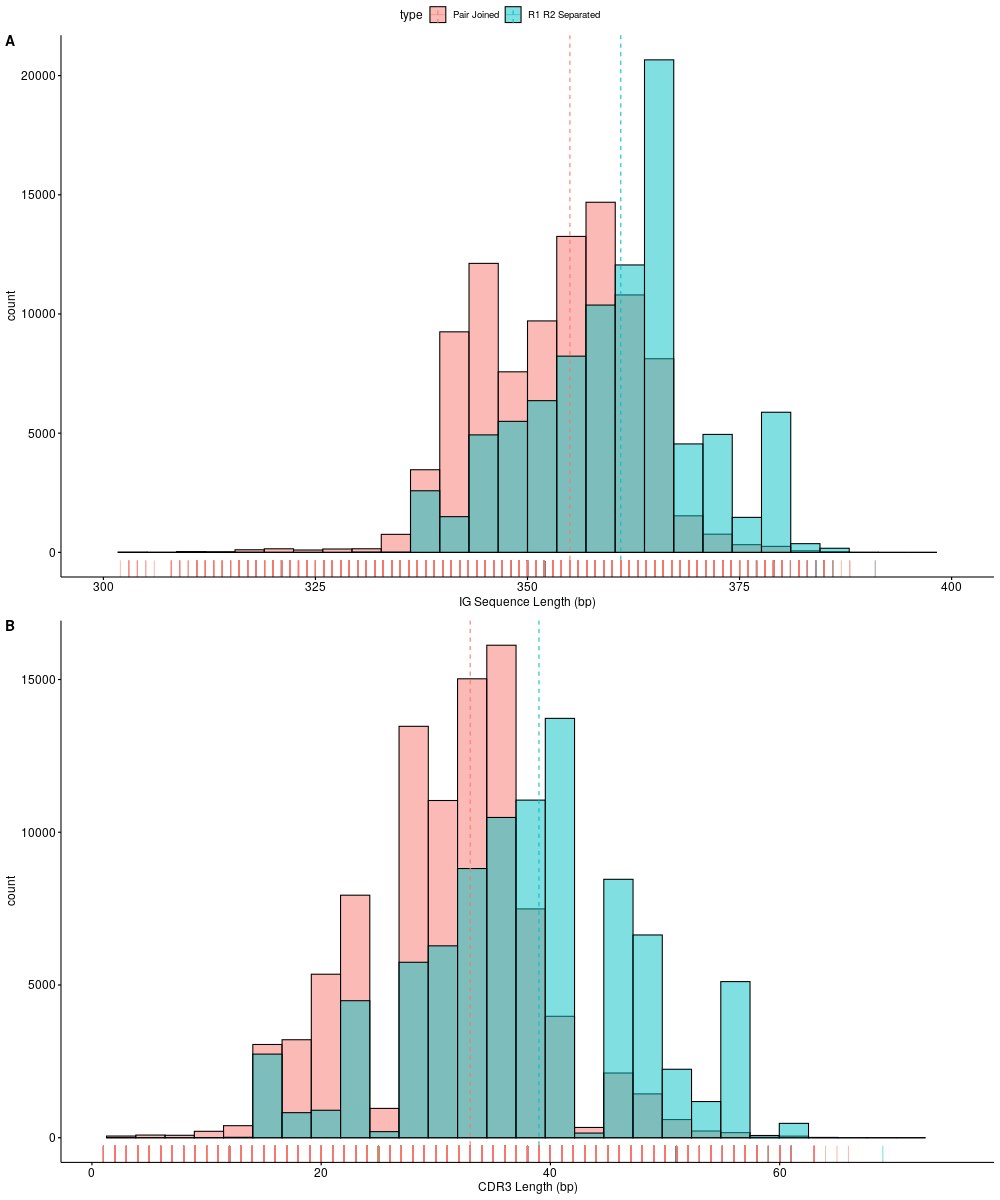

Figure S3. Pair-end joining results in biased sequence length and CDR3 length distributions. Pipelines with either pair-end joining or separated R1 and R2 processing were applied to same IgSeq data. The histograms of IG sequence length (A) and CDR3 length (B) were plotted by pooling all samples. Both CDR3 lengths and IG sequence lengths have shorter distributions after the pair-joining processing (red histograms) compared with the non-joining processing pipeline.

**Immunoglobulin gene filtering**

We relied on *bowtie2* (http://bowtie-bio.sourceforge.net/bowtie2/index.shtml, version 2.4.1) to filter out non-IG genes. Bowtie2 filtering was run with the following commands:

*bowtie2 -p 4 --local -D 20 -R 3 -N 1 -L 14 -i S,1,0.50 --mp 4,2 --no-unal*

*-x [IG gene database index] -1 [read 1 fastq file name] -2 [read 2 fastq file name]*

*--al-conc ./conc.fastq --un-conc ./unconc.fastq -S out.sam -X 600 -fr* .

The database index was built with IG gene sequences derived from IMGT and is available upon request. Two input files, read 1 and read 2 fastq files, were provided. The file, *out.sam*, is the output file that will be parsed and re-written as fastq files with only IG genes.

See <https://github.com/BULQI/IgSeq> for the scripts for IG filtering.

***Statistical Analysis*/ANOVA model**

We ran three-way ANOVA to test for the effects of three factors, tissue (spleen and bone marrow), isotype (IgM and IgG) and immunization (PBS, OVA, OVA+CpG and OVA+Alum). A linear statistical model with three-way interaction terms was used for the analysis,

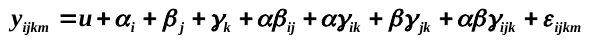

where
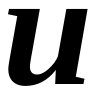
 is the grand mean;
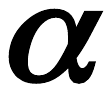
,
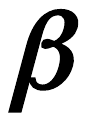
 and
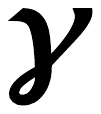
 are the main effects of the factor tissue, isotype and treatment, respectively;
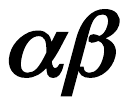
,
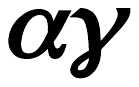
 and
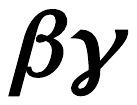
 are the two way interactions;
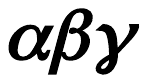
 is the three-way interaction term;
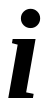
,
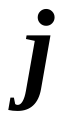
 and
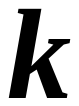
 are the level of different factors;
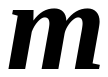
is the replicate number of each group. The model was fitted by linear regression and sum square errors were partitioned using the type II or type III ANOVA table. Analyses were carried out in R using the “car” package and post-hoc tests were done using the “emmean” package.

**IGHV Mutation Frequency Three-Way ANOVA**

Anova Table (Type III tests)

|  | Sum Sq | Df | F value | value | Pr(>F) |
| --- | --- | --- | --- | --- | --- |
| (Intercept) | 0.001205 | 1 | 468.8799 | <2.20E-16 | *** |
| tissue | 0.000106 | 1 | 41.1448 | 4.41E-07 | *** |
| treatment | 8.5E-07 | 3 | 0.1103 | 0.953444 |  |
| isotype | 0.000249 | 1 | 96.753 | 6.71E-11 | *** |
| tissue:treatment | 9.14E-06 | 3 | 1.1861 | 0.331651 |  |
| tissue:isotype | 9.5E-06 | 1 | 3.6965 | 0.064067 | . |
| treatment:isotype | 4.19E-05 | 3 | 5.4389 | 0.004154 | ** |
| tissue:treatment:isotype | 5.04E-06 | 3 | 0.6539 | 0.586798 |  |
| Residuals | 7.71E-05 | 30 |  |  |  |

A

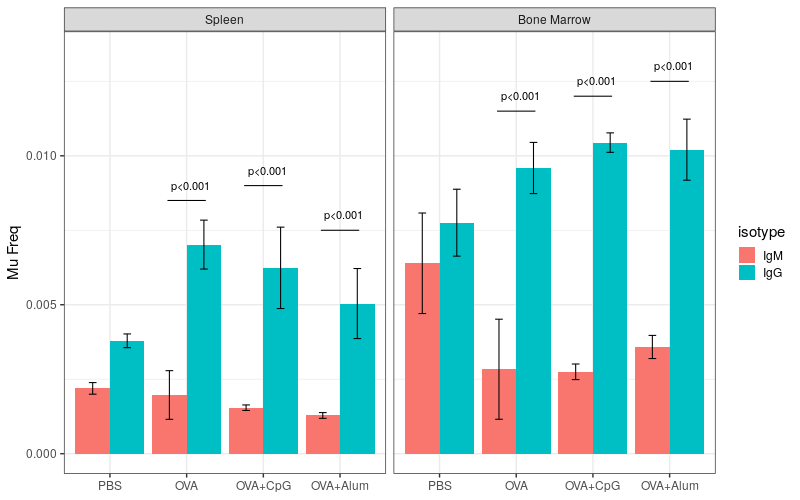

B

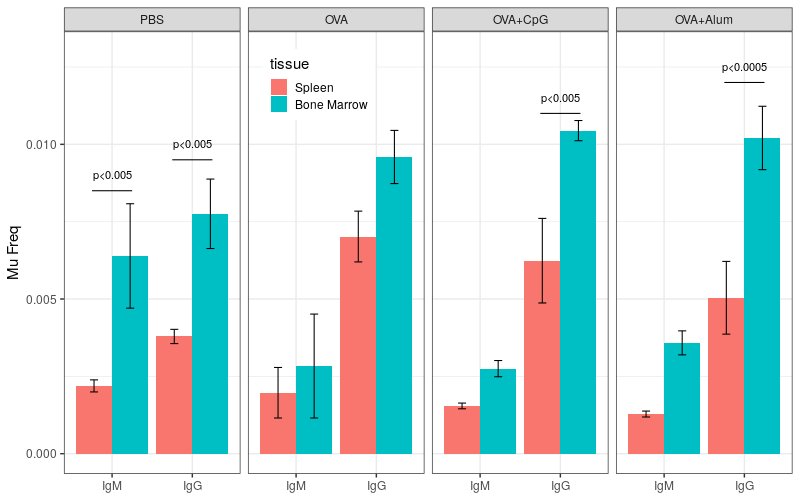

Figure S4. ~~IGHV mean mutation frequency difference between tissues. Mean mutation frequency difference between tissues was calculated by subtracting the mean mutation frequency of the spleen compartment from that of the bone marrow compart in each mouse by different isotypes. If, a difference with zero means equal mutation frequencies between different tissue compartments (the grey horizontal line), and a positive valued difference means the mutation frequency in the bone marrow compartment is higher. The differences were averaged in each group, and a line connected the compartments of the same isotype across different immunization group. (Blue line, the mean mutation frequency difference between bone marrow and spleen for IgM compartments; Red line, the difference between tissues for IgG compartments)~~. IGHV mean mutation frequency showed tissue and isotype differences. IGHV annotation was done by aligning to germline sequences with the software tool, Cloanalyst. Mutations were identified and expressed as per-nucleotide mutation frequency. The distributions of IGHV mutations, and means and standard deviations were obtained and used for statistical tests. A) IGHV mutation frequencies in different isotype compartments were compared in different tissue and under different immunizations. B) IGHV mutation frequencies in different tissues were compared under different immunizations by different isotypes. (Statistical p-values were corrected by FDR and significant p-values were indicated in figure)

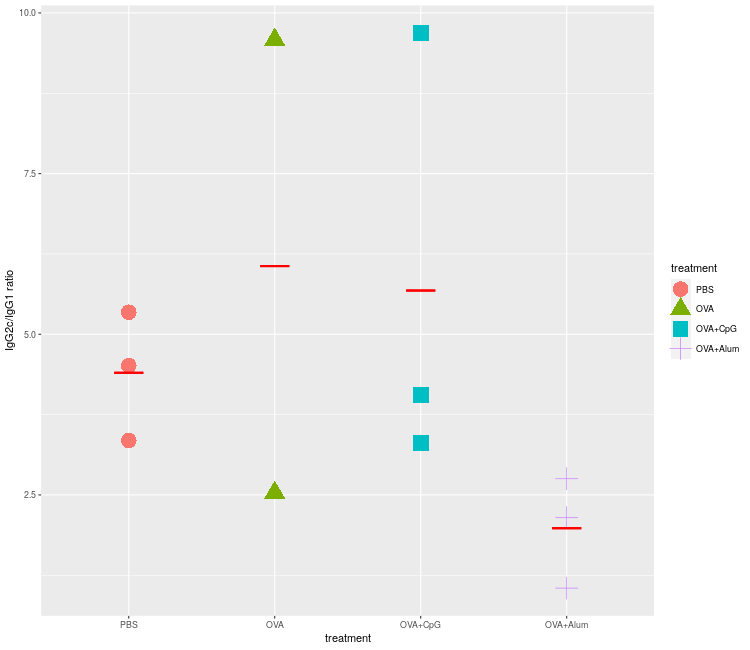

Figure S5. OVA immunization under different conditions (with or without adjuvants) changes the IgG subtype ratio. IgG subtypes were determined using the 5’- constant region sequence. The ratio of IgG2c/IgG1 were calculated for each sample. IgG subtypes have been shown to be valid indicators of preferential Th1/Th2 immune responses. A high IgG2c/IgG1 ratio usually associates with Th1 responses, and low IgG2c/IgG1 ratio Th2 responses. In this work, C57BL6J mice were used, which under the steady state (in PBS treated mice) showed a high IgG2c/IgG1 ratio. In OVA alone immunized and OVA+CpG treated mice, this ratio were slightly increased, respectively, although no statistical significance was achieved. While OVA+Alum immunization resulted in decreased ratio of IgG2c/IgG1, indicating a biased response towards Th2.

**Immunoglobulin Heavy Chain Variable Gene Usage:**

*Compositional Data Transformation and Principal Component Analysis (PCA)*

In order to analyze heavy chain variable gene usage (IGHV) data, we first transformed gene usage counts into compositional data using the R compositions library. In the next step, the data were further undergone the center log ratio (CLR) conversion, which transforms/maps the Aitchison simplex into the real space. Lastly, PCA was carried out on the CLR transformed data with the hypothesis that some of the obtained PCs might reflect the biological differences caused by treatment effects.

As a result, the first 6 PCs capture 58% of total contribution, 15.3%, 13.0%, 9.5%, 7.3%, 6.8%, 5.8%, and were chosen for the further ANOVA analysis for testing the treatment effects.

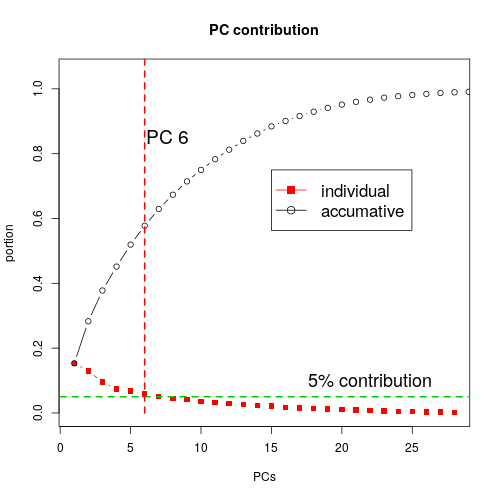

Figure S6. The accumulative and individual PC contributions. The contribution is the percentage of variation a specific PC explains. The first six PCs were chosen, since they together explain about 58% of the total variation and each of them has a larger than 5% contribution.

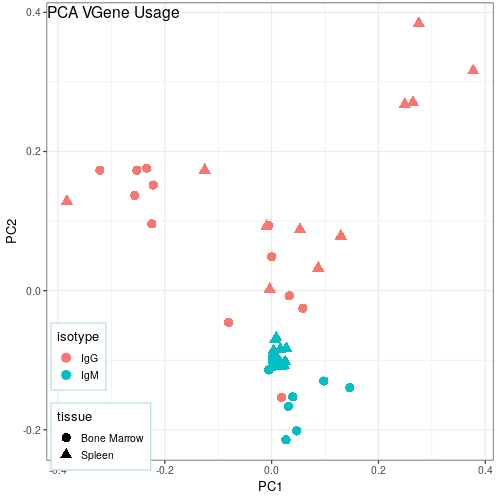

Figure S7. The projection of IGHV gene usage data on the PC1 and PC2 coordinates. Data points are colored by isotypes and shaped by tissues.

***Three-way ANOVA for IGHV gene usage***

After PCA, samples are associated with PC scores, which are the new variables obtained by taking the inner-product between the corresponding eigen vector and the scaled gene usage compositional proportion. The PC score can been seen as a reflection of underlying treatment effects. The PC scores were used to fit for a linear three-way model (tissue, isotype and immunization) for ANOVA. In the case there are significant effects in ANOVA, “Pr(>F)” <0.05, follow-up tests are done accordingly with linear contrasts. The results of ANOVA and follow-up tests are shown below.

PC1 (Tissue and treatment effects)

*ANOVA (Type II)*

|  | Sum Sq | Df | F value | Pr(>F) |  |
| --- | --- | --- | --- | --- | --- |
| Treatment | 68.964 | 3 | 3.1891 | 0.037784 | * |
| Isotype | 20.521 | 1 | 2.8469 | 0.101923 |  |
| Tissue | 70.136 | 1 | 9.73 | 0.003984 | ** |
| immunization:isotype | 25.584 | 3 | 1.1831 | 0.33273 |  |
| immuniztion:tissue | 113.119 | 3 | 5.231 | 0.005037 | ** |
| isotype:tissue | 103.076 | 1 | 14.2998 | 0.000694 | *** |
| immunization:isotype:tissue | 81.191 | 3 | 3.7545 | 0.021136 | * |
| Residuals | 216.247 | 30 |  |  |  |

PC2 (Isotype effects)

|  | Sum Sq | Df | F value | Pr(>F) |  |
| --- | --- | --- | --- | --- | --- |
| immunization | 30.08 | 3 | 4.796 | 0.007591 | ** |
| isotype | 372.25 | 1 | 178.0617 | 3.76E-14 | *** |
| tissue | 31.44 | 1 | 15.0386 | 0.000533 | *** |
| immunization:isotype | 9.63 | 3 | 1.5354 | 0.225622 |  |
| immunization:tissue | 45 | 3 | 7.1752 | 0.000903 | *** |
| isotype:tissue | 4.5 | 1 | 2.1521 | 0.15278 |  |
| immunization:isotype:tissue | 35.33 | 3 | 5.6327 | 0.003478 | ** |
| Residuals | 62.72 | 30 |  |  |  |

PC3 (immunization and isotype effects)

|  | Sum Sq | Df | F value | Pr(>F) |  |
| --- | --- | --- | --- | --- | --- |
| immunization | 134.349 | 3 | 6.4537 | 0.001674 | ** |
| isotype | 0.063 | 1 | 0.0091 | 0.924522 |  |
| tissue | 4.54 | 1 | 0.6542 | 0.424967 |  |
| immunization:isotype | 34.604 | 3 | 1.6623 | 0.196115 |  |
| immunization:tissue | 2.256 | 3 | 0.1084 | 0.954546 |  |
| isotype:tissue | 9.765 | 1 | 1.4072 | 0.244824 |  |
| immunization:isotype:tissue | 42.205 | 3 | 2.0274 | 0.131201 |  |
| Residuals | 208.174 | 30 |  |  |  |

Contribution of variables to PCs:

There are two kinds of contributions in PCA. The first one is on how each PC is made up by different gene usages. This is quantified by the relative weight of each gene usages in the PC, which is the corresponding coefficient of the eigen vector of the PC. To obtain the percentage contribution, we need to compute the sum square percentage of each eigen vector. The second is about how each gene usage contribute to different PCs. This contribution is obtained by comparing the relative contributions of the same gene usage to different PCs. Moreover, the sign of the coefficients in the eigen vector defines the direction of the change of gene usages relative to the PC.

PC1:

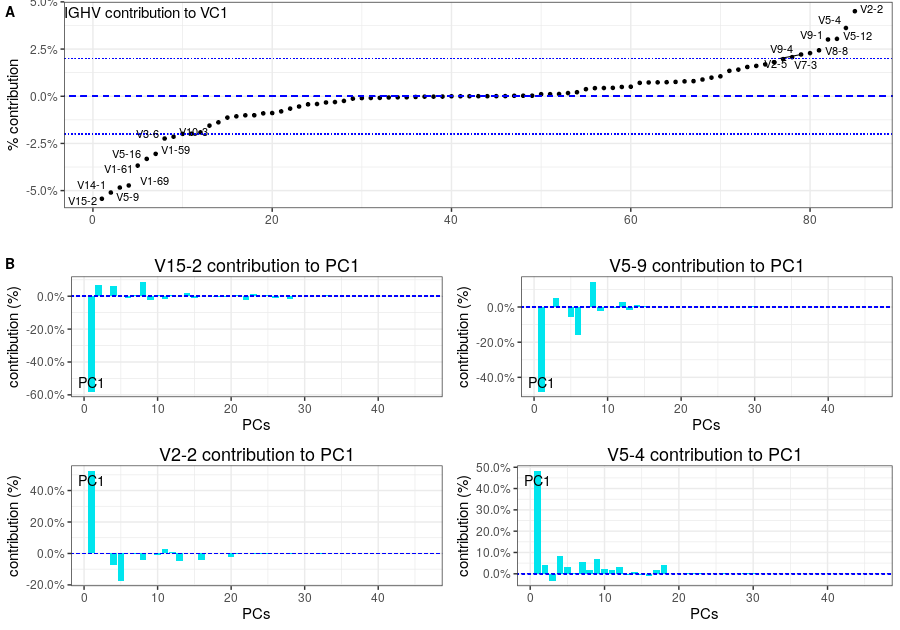

Figure S8. Contribution of IGHV gene usage to PC1. Two types of contributions are plotted. (A) The contribution of gene usage to PC1. The IGHV gene segments that contribute more than 2% of total variation are labelled. (B) The top two contributing gene segments (of both negative and positive signs) were shown individually for their contributions to all PC.

PC2

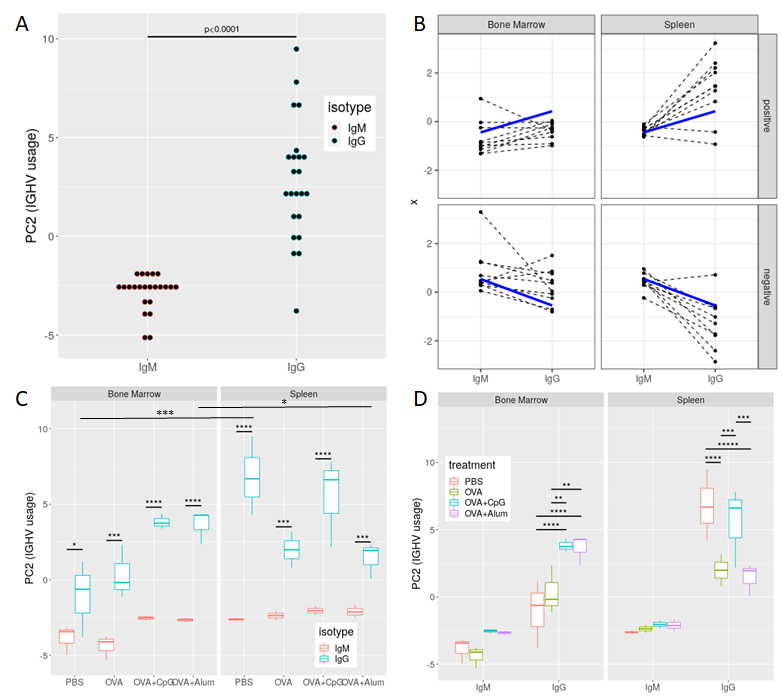

Figure S9. PC2 of IGHV gene segment usage showed significant isotype difference. IGHV gene segment usage were determine and transformed into compositional data format. PCA were carried out on the compositional usage and top six contribution PCs were picked for further statistical analysis. A) Distribution of PC2 by isotype. B) Raw gene usages before PCA analysis were plotting to verify the pattern (for both positively and negatively changed genes). C) Comparison of PC2 between isotypes by tissue and treatment. D) Comparison of PC2 between isotypes by tissue and isotype. (*, p<0.05; **, p<0.01; ***, p<0.005; ****, p<0.001).

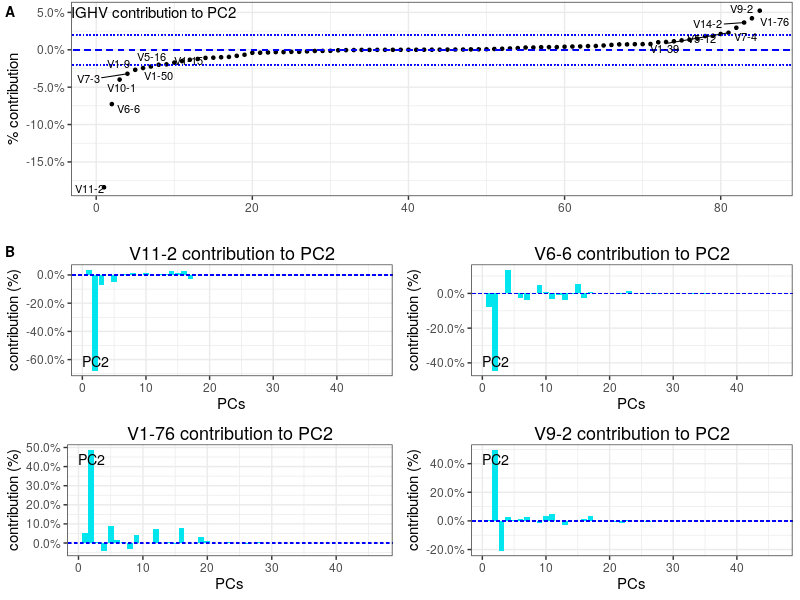

Figure S10 Contribution of IGHV gene usage to PC2. Two types of contributions are plotted. (A) The contribution of gene usage to PC1. The IGHV gene segments that contribute more than 2% of total variation are labelled. (B) The top two contributing gene segments (of both negative and positive signs) were shown individually for their contributions to all PC.

PC3

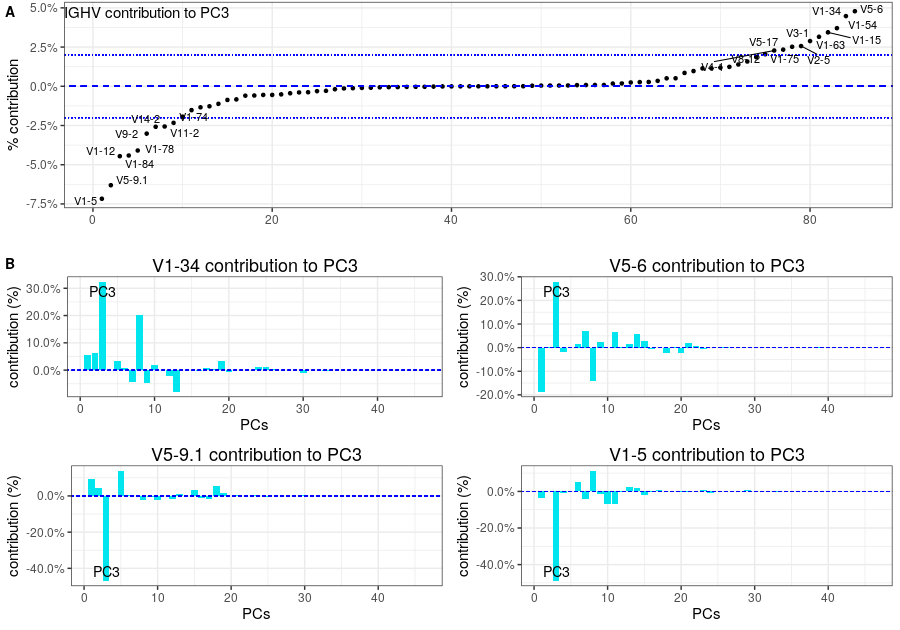

Figure S11 Contribution of IGHV gene usage to PC3. Two types of contributions are plotted. (A) The contribution of gene usage to PC3. The IGHV gene segments that contribute more than 2% of total variation are labelled. (B) The top two contributing gene segments (of both negative and positive signs) were shown individually for their contributions to all PC.

**IGH CDR3 Lengths**

| A | 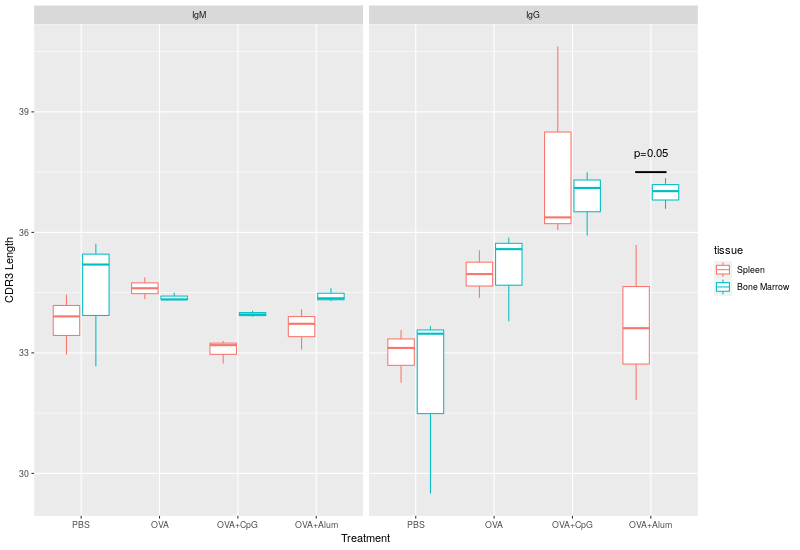 |
| --- | --- |
| B | 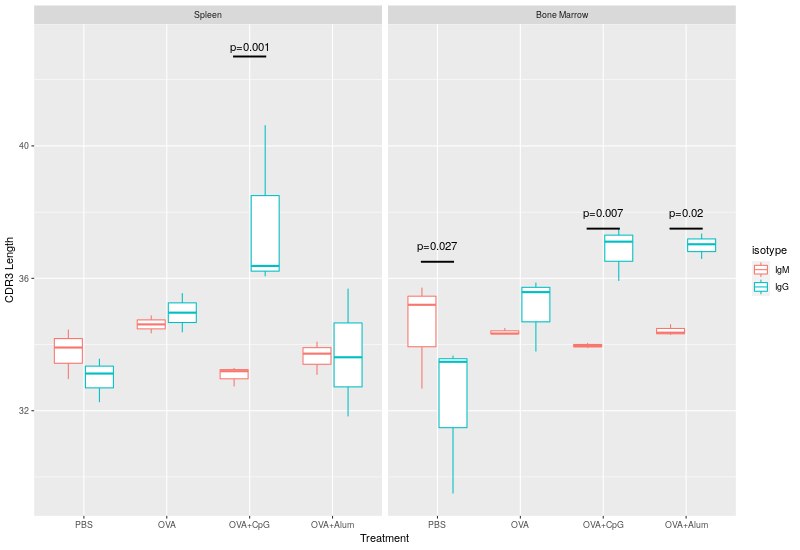 |

Figure S12 IGHV CDR3 lengths were affected by OVA immunizations with adjuvants. The distribution of CDR3 lengths was determined for each sample and mean CDR3 lengths were calculated. Three-way ANOVA was carried out to identify factor effects, including tissue, isotype and immunization. The results showed a significant isotype and immunization interaction effect, p< 1.0e^-5^. Here we plot to show differences between tissues (A) and isotypes by different immunizations (B). (The p-values are the results of the post hoc tests by linear contrasts and corrected by FDR).

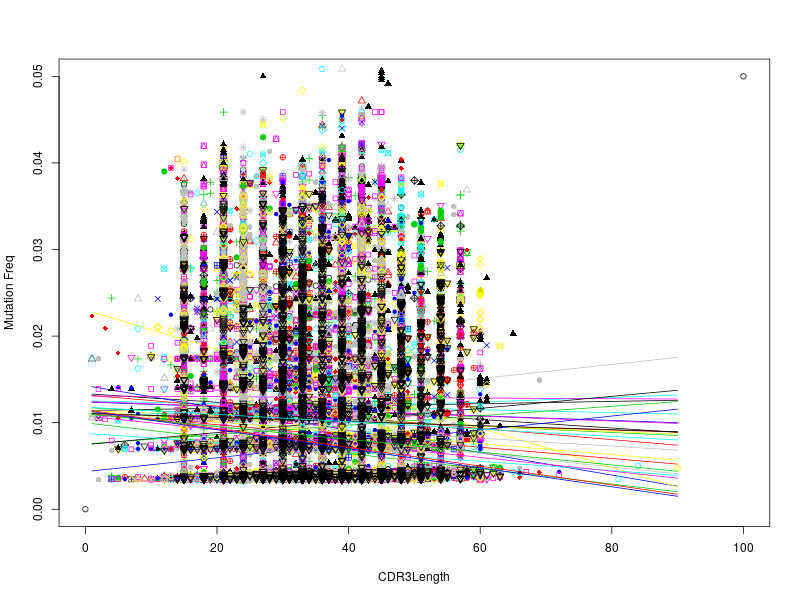

Figure S13. IGHV CDR3 length did not correlate with sequence mutation frequency. The CDR3 length and mutation frequency was plotted by each sample. Linear regressions were done to fit straight lines between them. Correlation coefficients between CDR3 length and mutation frequency were also calculated, and all had values close to zero (between -0.18 and 0.11).

**PBS	OVA**

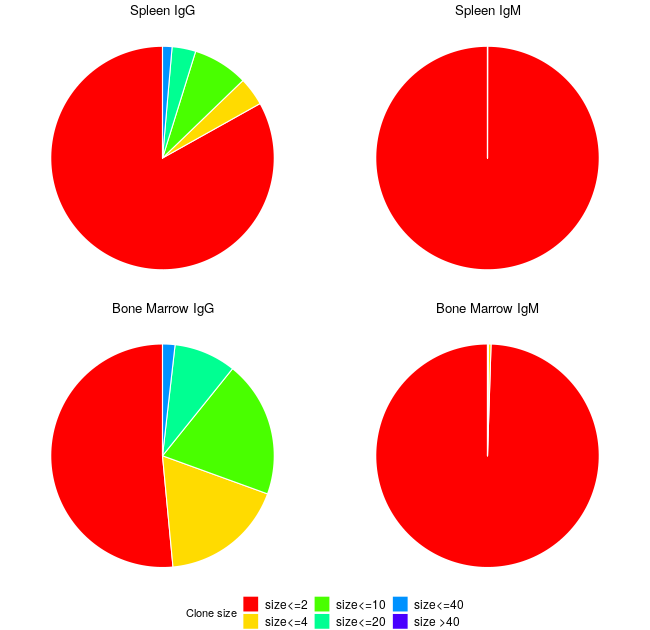

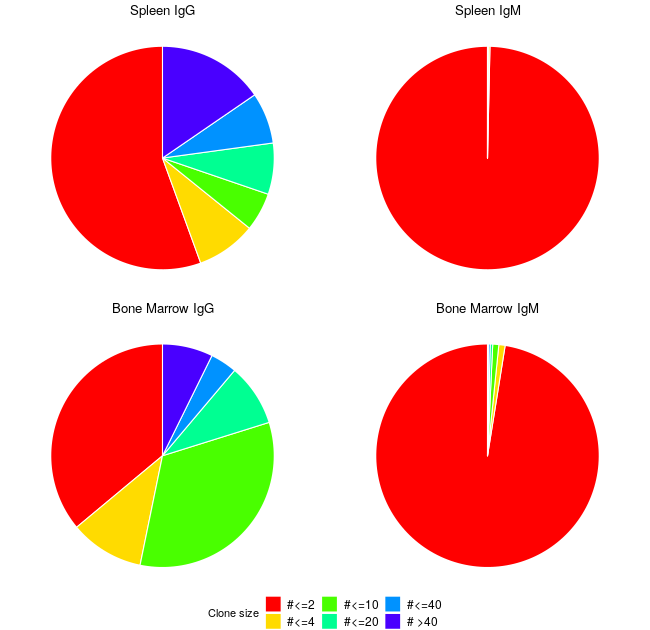

**OVA+CpG**

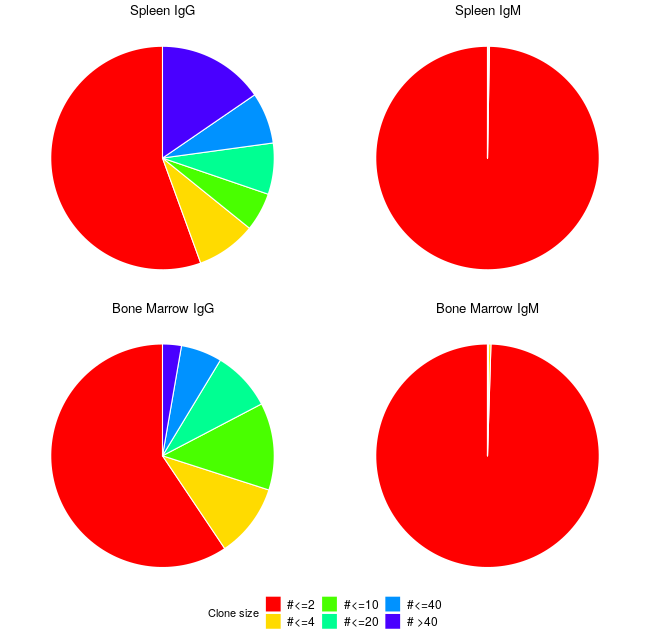

Figure S14. The distributions of clone sizes in different tissue and isotype compartments. Clones were grouped by their size as indicated by the legend. Example clone distributions are shown for different immunization groups. In IgM compartments, the majority of clones are small clones (size of 1 or 2 members) and in the memory repertoire (IgG), the larger clones are almost 50% of the total. The similar patterns are observed in both bone marrow and spleen.

**Diversity (HILL numbers):**

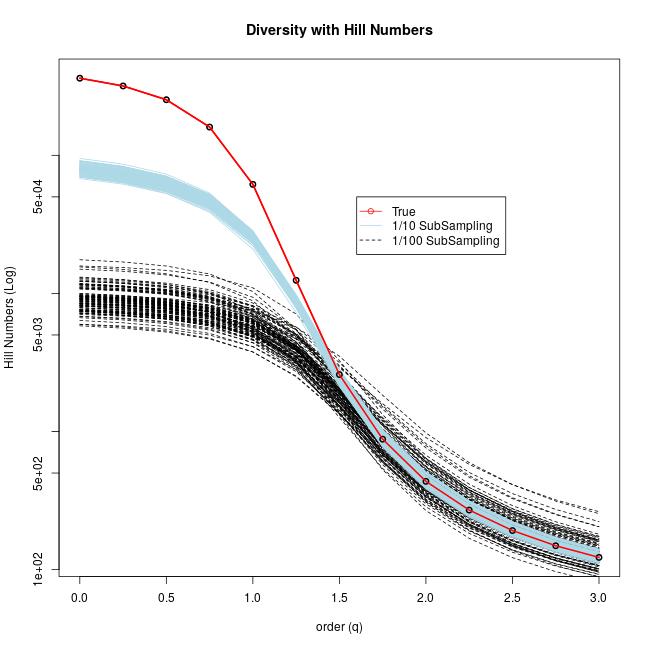

Figure S15. The effect of sampling depth on diversity profiles. It is known that incomplete sampling is a major cause for bias in estimating the diversity profile of low orders. In order to determine the effect of sequencing depth on our clonal diversity profiles, we ran subsampling of the sequencing data to different levels. Here we showed the result of one such run on one sequencing sample. The true diversity profile (red line) was obtained with all the sequences in the sample. Then we run subsampling randomly with 1/10 (light blue line) and 1/100 (black dashed line) of the total number of sequence to estimate the diversity profile. The subsampling was repeated 100 times. We can see in our results that incomplete sampling affected the diversity profile estimation of orders less than 1.5.

A

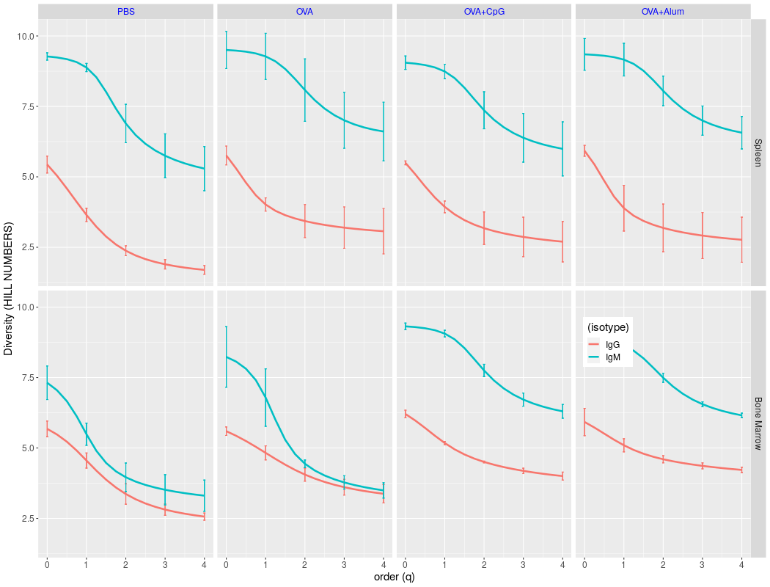

B

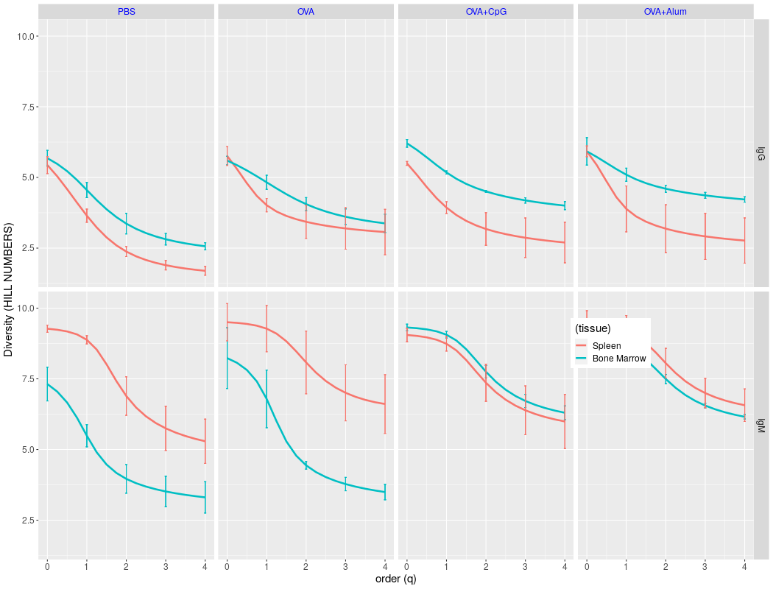

C

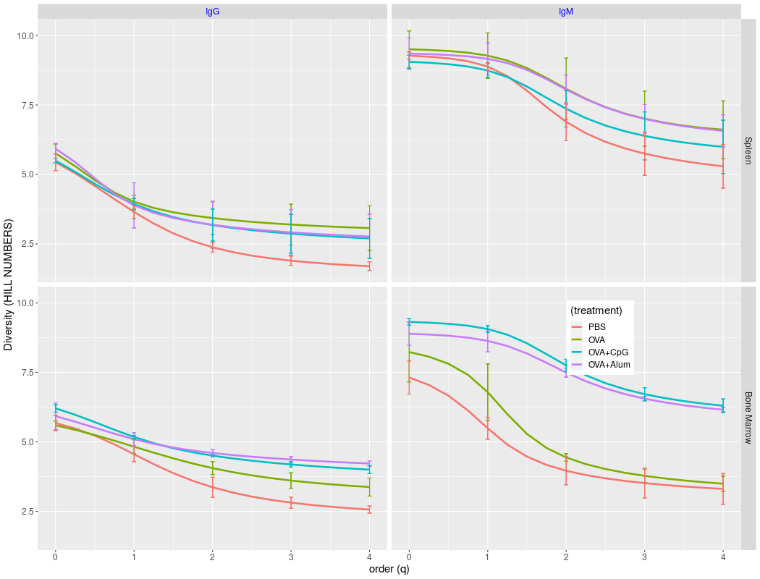

Figure S16. Clonal diversity expressed as Hill numbers. Ig sequences were annotated and partitioned into clones. Then the diversity profile was expressed as Hill numbers with orders between 0 and 4. A) Comparison of diversity profiles by isotypes. Each line is for one isotype, and sub-plots are arranged for different tissues and immunization status. Standard errors are plotted at the order of 0, 1, 2, 3 and 4. B) Comparison of diversity profiles by tissues. Each line is for one tissue, and sub-plots are arranged for different isotype compartments and immunization status. Standard errors are plotted at the order of 0, 1, 2, 3 and 4. C) Comparison of diversity profiles by immunizations. Each line is for one immunization status, and sub-plots are arranged for different isotypes and tissues. Standard errors are plotted at the order of 0, 1, 2, 3 and 4.

A

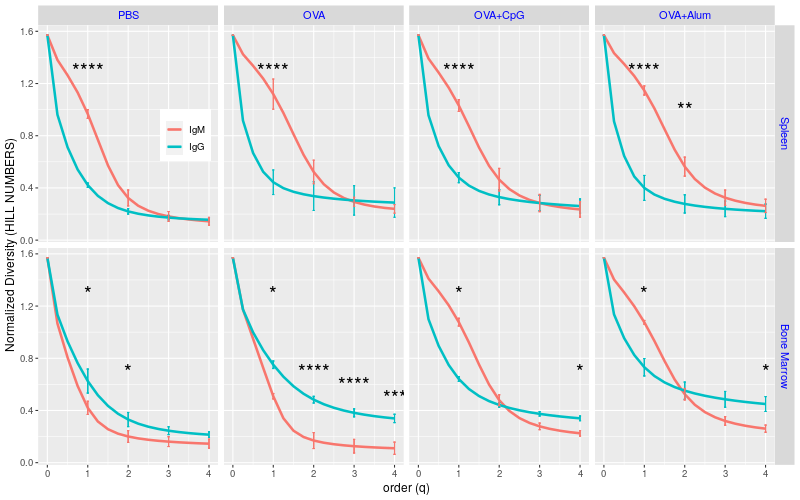

B

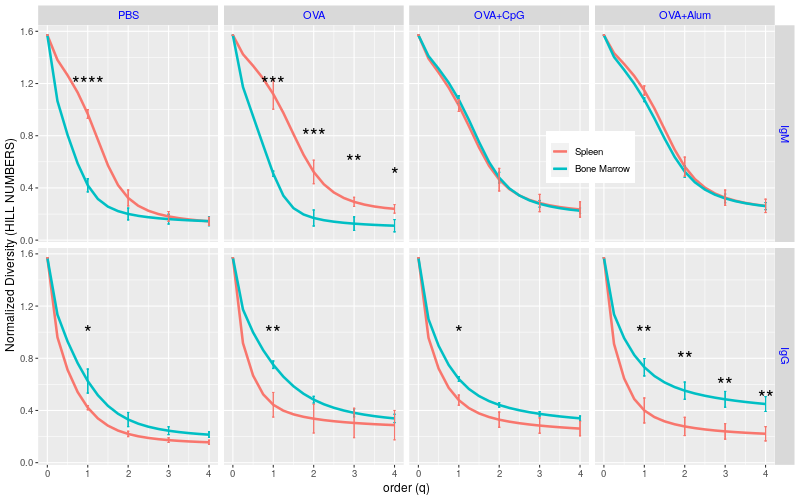

Figure S17. Normalized/re-scaled clonal diversity profiles. Ig sequences were annotated and partitioned into clones. Then the diversity profile was first expressed as Hill numbers with orders between 0 and 4 and then was normalized/re-scaled with the clone richness. A) Comparison of diversity profiles by isotypes. Each line is for one isotype, and sub-plots are arranged for different tissues and immunization status. Standard errors are plotted at the order of 0, 1, 2, 3 and 4. B) Comparison of diversity profiles by tissues. Each line is for one tissue, and sub-plots are arranged for different isotype compartments and immunization status. Standard errors are plotted at the order of 0, 1, 2, 3 and 4.

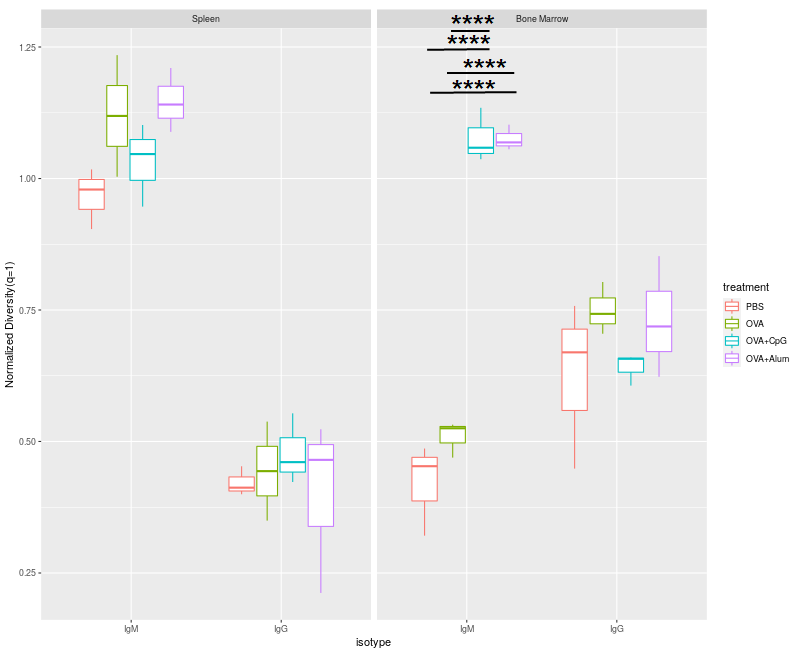

Figure S18. Immunization effects on clonal diversities (order q=1). Clonal diversity profiles were determined on clone abundance data. The profiles were further normalized by clonal richness. ANOVAs were run to identify significant factor effects at each order of diversities, and follow-up tests were used to find specific group differences. Here we compared diversities (q=1) of clone samples under different immunization conditions in different tissue and isotype compartments. (****, p<0.001).

**Intra-clonal diversity / Clonal selection**

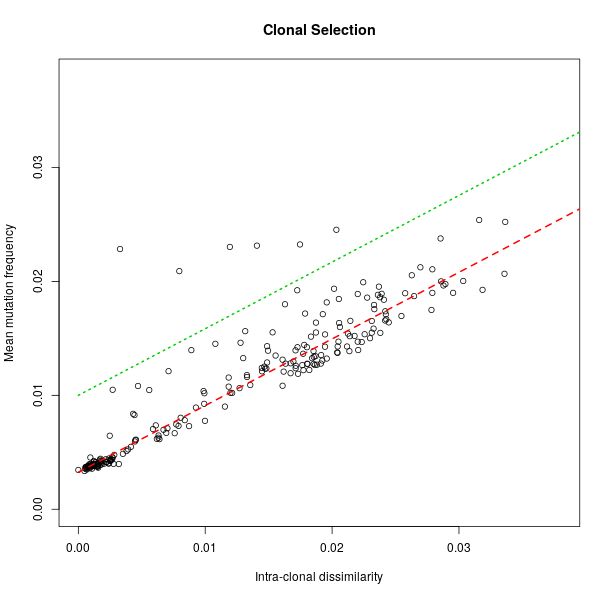

Figure S19. The clonal selection positive gate (threshold). Largest clones from PBS IgM compartments of three mice were pooled and used to set the negative gate/threshold for clones undergone selection. Intra-clonal sequence dissimilarity and mean mutation frequency of these clones were plotted on a 2-D x-y figure. Most clones show no signs of selection, growing proportionally the dissimilarity and mean mutation frequency, and locate closely to a straight line (the red dashed line, r=0.96). We draw a threshold line (the green dotted line), the points above which are clones enriched with members be selected in favor and ones below which are clones showing no obvious selections.

| A | 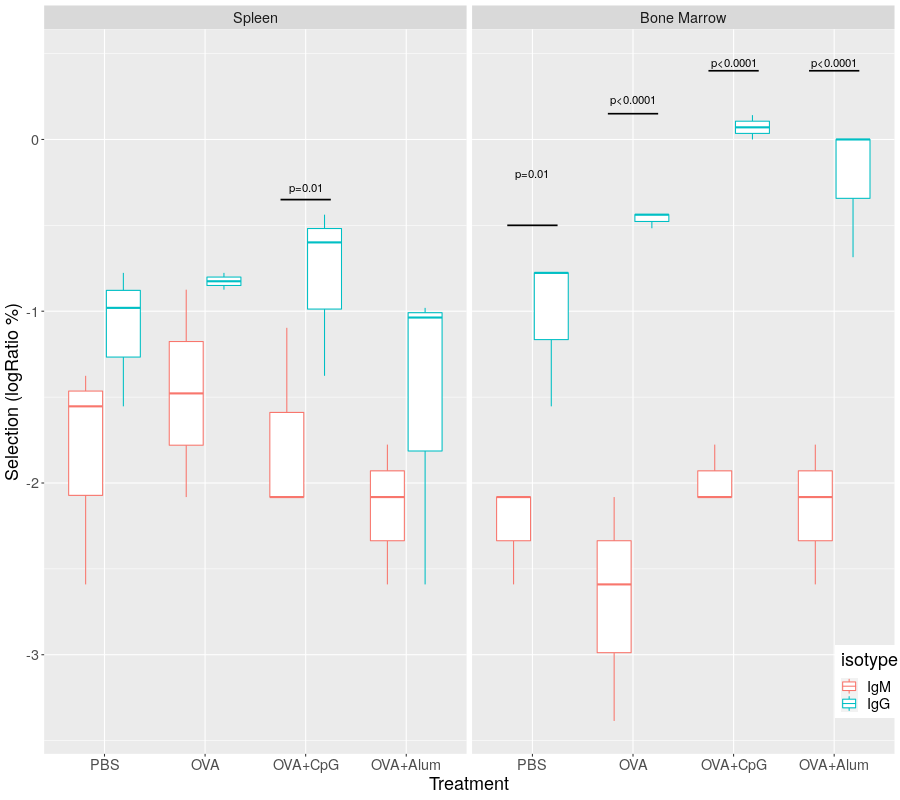 |
| --- | --- |
| B | 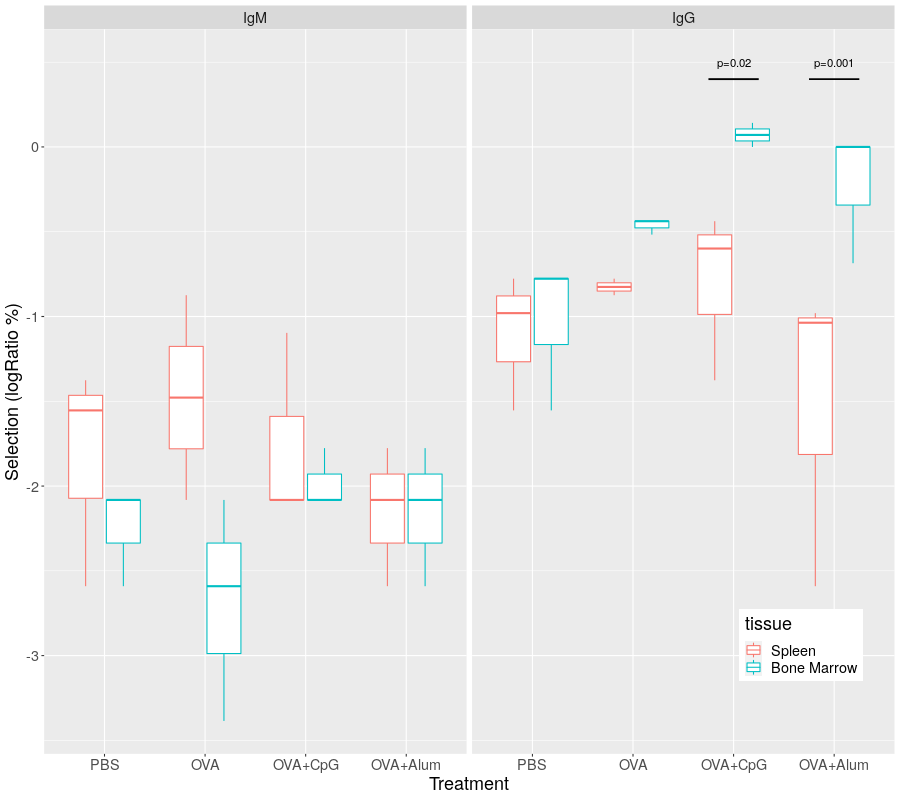 |

Figure S20. Distribution and comparison of percentages of clones showing signs of being selected in different tissues and isotypes. Top clones with largest size in different compartments were evaluated based on two parameters, mean mutation frequency and clonal pairwise similarity. The threshold/gate for clones being selected was set up using the spleen IgM clone data showing on the two dimension plot (Fig. S19). Clones locating above the threshold line was treated as the ones being selected. The percentage of clones being selected in the top clones in each sample was plotted and analyzed for factor effects. A) Distribution and comparison of percentages of clones being selected in different isotypes. B) Distribution and comparison of percentages of clones being selected in different tissues. (Significant p values were indicated).

**Unsupervised machine learning (Multiple Factor Aanlysis)**

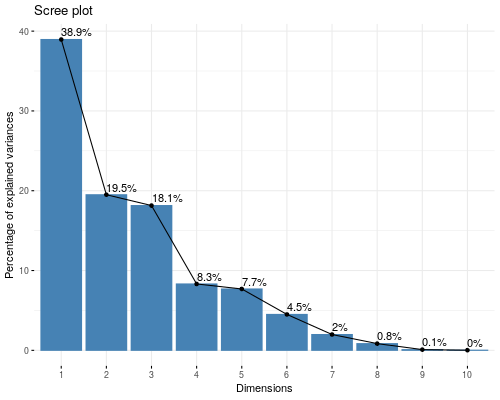

Figure S21. Percentage contribution of top 10 principal components (PCs). Multiple factor analysis (MFA) was carried out on the data pooled together with groups of variables including gene usage, mutation frequency, CDR3 length, clone diversity (expansion) and clonal selection level. The percentage contributions of top 10 PCs (dimensions) were shown and the first three PCs explained combined ~77% of total variation.

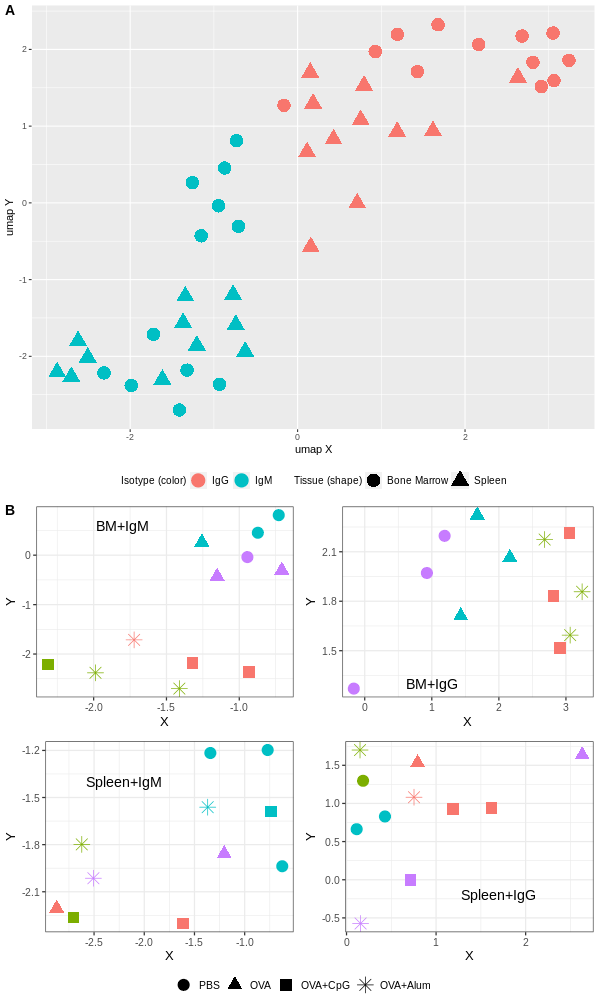

Figure S22 Un-supervised clustering of the Ig repertoire data. We ran unsupervised clustering on the MFA projected data with all PCs. Different clustering algorithms were applied and similar results were obtained. Here we show the spectral clustering results. Data were first visualized on UMAP-transformed 2D coordinates by their isotypes (color) and tissues (shape) (A). Then, the clustering results generated by the spectral clustering algorithm were showed in different compartments. In each compartment, samples receiving different immunizations were shown in different shapes and the color of data points indicated the clusters they belonged to.

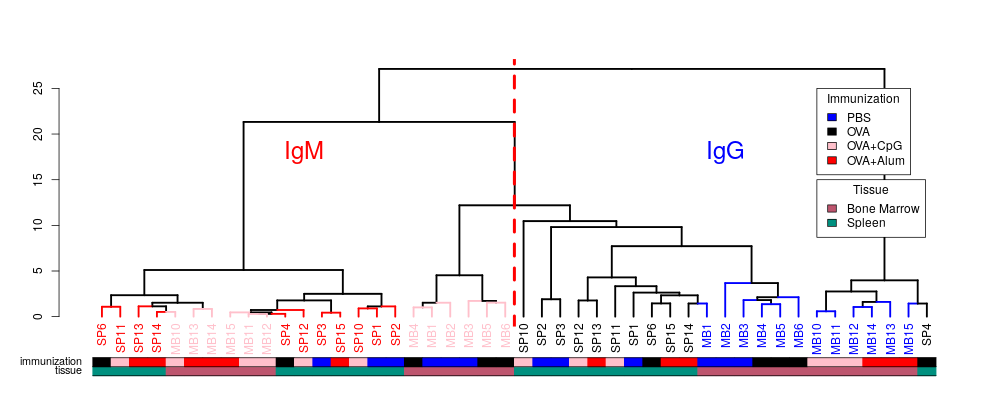

Figure S23. Hierarchical clustering of Ig sequencing data. MFA was carried out on groups of variables including V gene usage, mutation frequency, CDR3 length, clone diversity (expansion) and clone selection strength. The projected data on the new PCs were then used for hierarchical clustering to find signature changes by different factors. We could see that data from different isotypes are well separated. Within each isotype, data points from different tissues are also largely separated. However, data from different immunization conditions are arranged into groups in bone marrow compartments, especially in the bone marrow IgG compartment. In spleen compartments, data points from different immunization groups are mixed.

See attached supplementary file movie1.gif

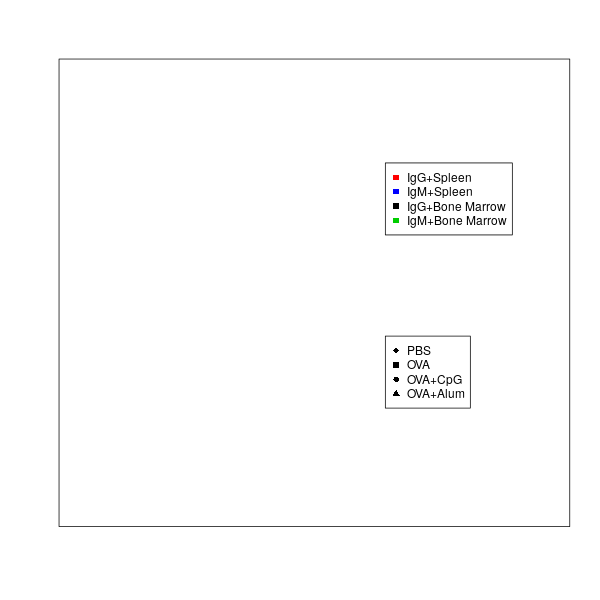

Figure S24. Projection of individual data on the first three PC space of the MFA analysis. MFA was carried out on groups of variables including V gene usage, mutation frequency, CDR3 length, clone diversity (expansion) and clone selection level. Individual observations were then projected on the first three PC coordinates to show the correlations between them. Observations of different isotypes stay at two well separated locations. Within the same isotype, observations of different tissues also form largely separated clusters. Furthermore, the data points of the IgG isotype show large variations, and the IgM data show little. A 3-D rotating gif picture was saved as a separate file, named “movie1.gif”. Here we show the figure legends. Different compartments are in different color: Red, IgG spleen; black, IgG bone marrow; green, IgM bone marrow; blue, IgM spleen. Immunization conditions are in different shape: diamond, PBS; square, OVA; circle, OVA+CpG; triangle, OVA+Alum.

See attached supplementary file “movie2.gif”.

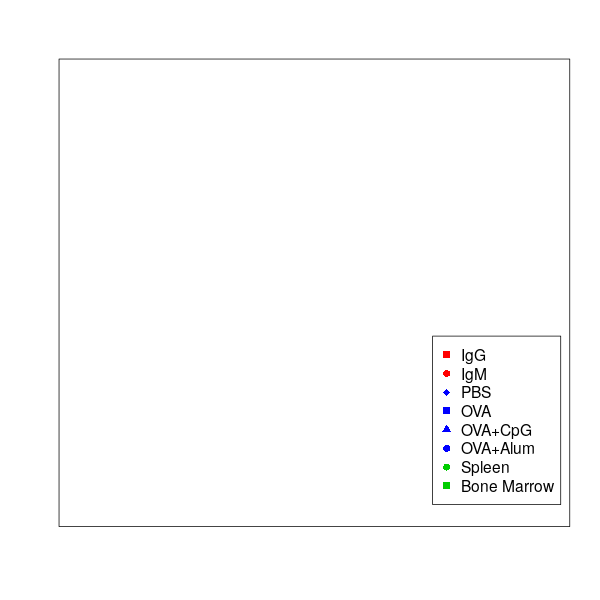

Figure S25. Correlation and projection of supplementary (grouping) variables on the first three PC dimensions. MFA was carried out on groups of variables including V gene usage, mutation frequency, CDR3 length, clone diversity (expansion) and clone selection level. Supplementary variables such as isotype, tissue and immunization condition were also included. These variables were then projected on the first three PC planes to show the correlations between them. Isotype IgM and IgG are distantly separated mainly along the first PC. Tissue bone marrow and spleen spread almost orthogonally to isotype and locate roughly along the second PC. Treatment variables correlate with tissue variables but less with isotype variables. The PBS control and CpG groups stay close to spleen, the Alum group locates next to bone marrow, and the OVA alone positions near the origin. Furthermore, the PBS and OVA groups stay closer to the IgM isotype, but two adjuvant groups locate near the IgG. A 3-D rotating gif picture was saved as a separate file, named “movie2.gif”. Here we show the figure legends. Different grouping variables are in different color: red, isotype; green, tissue; blue, immunization. Within each group, different point shapes indicate different categories.

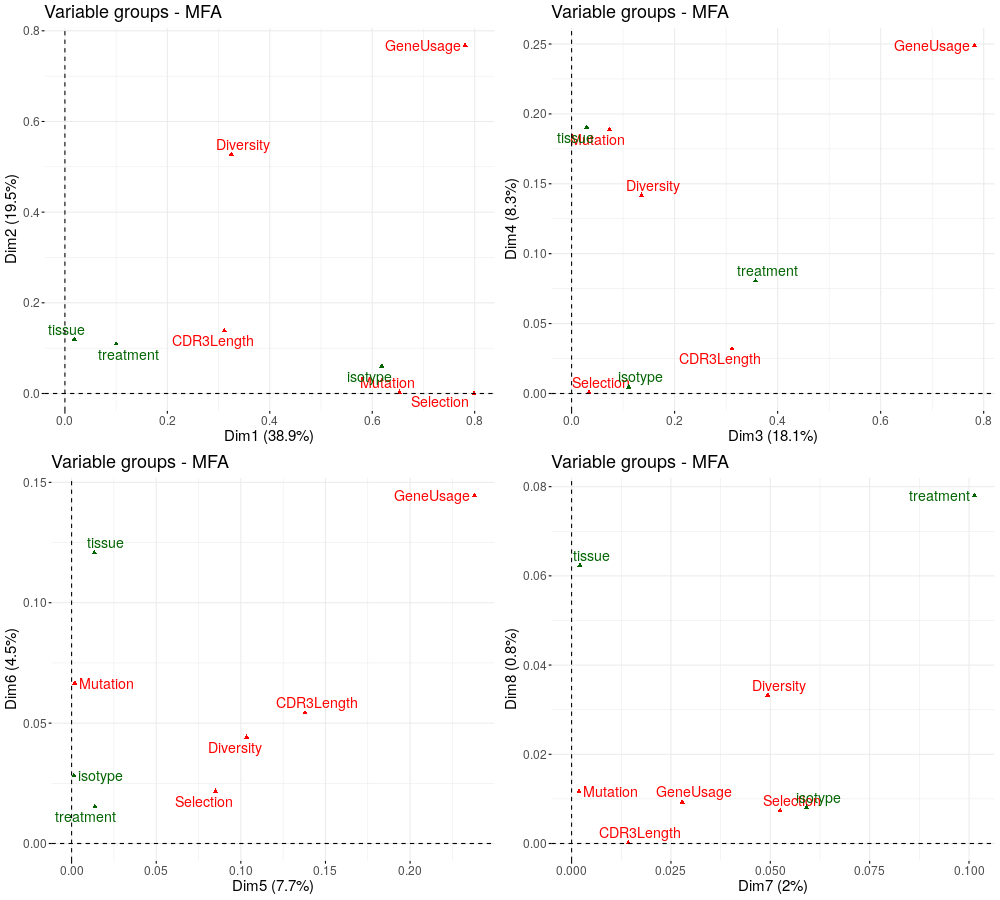

Figure S26. Correlation and projection of variables on new PC coordinates. MFA was carried out on groups of variables including V gene usage, mutation frequency, CDR3 length, clonal diversity (clonal expansion) and clone selection level, which are labelled as red texts. Supplementary variables or grouping variables, such as isotype, tissue and treatment were also included as green labels. These variables were then projected on the new PC coordinates to show the correlations between them. PC1 (labeled as Dim1 in the figure) is mainly explained by the variations caused by different isotypes, PC2 (labeled as Dim2 in the figure) is more about combined treatment and tissue variations, PC3 (labeled as Dim3 in the figure) is about the treatment difference, and PC4 (labeled as Dim4 in the figure) are for variations due to different tissues.

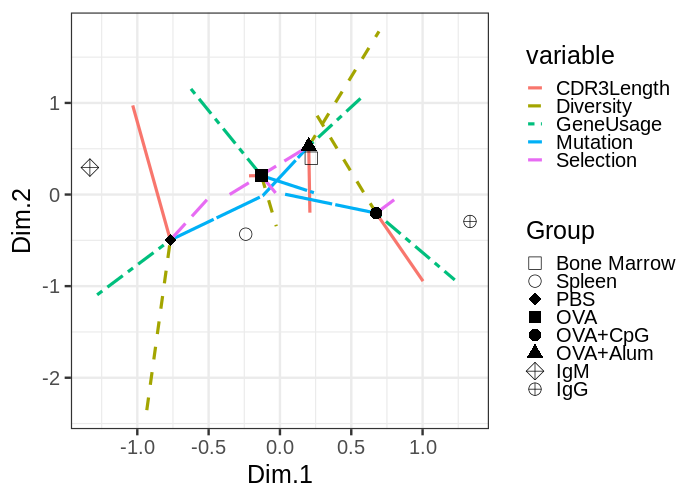

Figure S27. The partial group graph of supplementary category variables in MFA. In this graph, supplementary categorical variables (such as isotype, tissue and immunization), as a result of averaging data of their member samples, was plotted. The partial group points are the results obtained from the analysis performed with a single variable (or a single variable group, in this case, gene usage, mutation, CD3 length, clone diversity and clone selection level). In other words, this is the information of a category considered from the point of view of a single variable group. Supplementary category variables were plotted as points with different shapes at the barycenter, to which variable groups were connected by lines with different color. The figure was arranged into sub-plots by isotype, tissue (see Fig 7G in the main text) and immunization for easy visualization. (Shape: open symbols for isotypes; filled one for tissues; crossed ones for immunizations. Line type: solid line for CDR3 length; dashed line for Diversity; dotted dash line for gene usage; long dash for mutation frequency; two dash for selectin level).

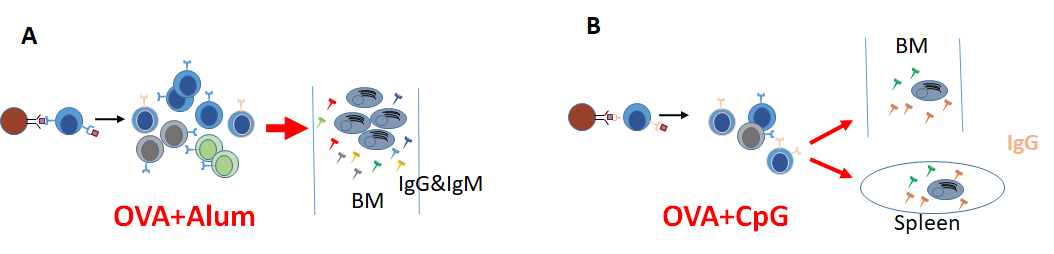

Figure S28. Proposed models for Alum and CpG effects on Ig repertoire in OVA immunized murine model. A) The Alum adjuvant in OVA immunization results in increased clone diversity with higher effective number of large clones and mainly impacts the BM compartment of both IgG and IgM isotypes (a quantity model). B) The CpG adjuvant leads to a higher degree of clonal selection and affects repertoires in both spleen and BM compartments of mainly IgG isotype (a quality model).
