## Supplementary figures and images for "Characterizing adjuvants’ effects at the murine immunoglobulin repertoire level"

### Supplementary Material File 2

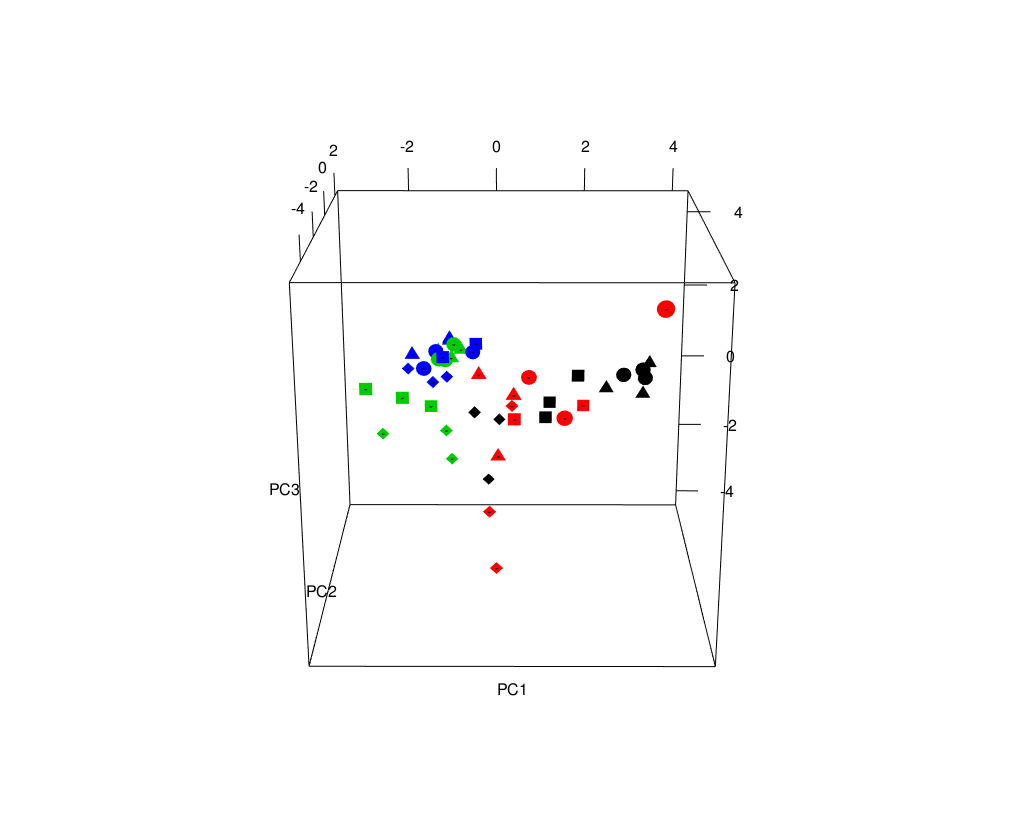

### Supplementary Material File 3

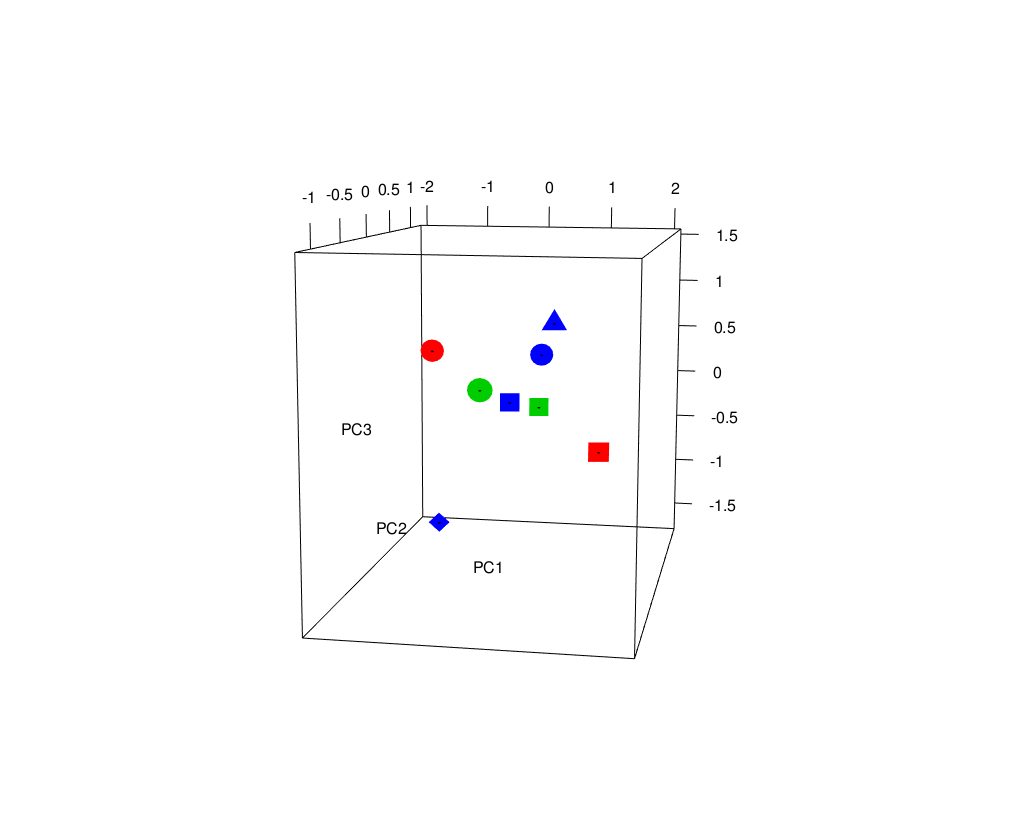
